# State and stimulus dependence reconcile motion computation and the *Drosophila* connectome

**DOI:** 10.1101/2021.04.17.440267

**Authors:** Jessica R. Kohn, Jacob P. Portes, Matthias P. Christenson, LF Abbott, Rudy Behnia

**Author notes:** equal contribution (Jessica R. Kohn, Jacob P. Portes).

## Abstract

Sensory systems dynamically optimize their processing properties in order to process a wide range of environmental and behavioral conditions. However, attempts to infer the function of these systems via modeling often treat system components as having static processing properties. This is particularly evident in the *Drosophila* motion detection circuit, where the core algorithm for motion detection is still debated, and where inputs to motion detecting neurons remain underdescribed. Using whole-cell patch clamp electrophysiology, we measured the state- and stimulus-dependent filtering properties of inputs to the OFF motion-detecting T5 cell in *Drosophila*. Simply summing these inputs within the framework of a connectomic-constrained model of the circuit demonstrates that changes in the shape of input temporal filters are sufficient to explain conflicting theories of T5 function. Therefore, with our measurements and our model, we reconcile motion computation with the anatomy of the circuit.

## Introduction

Flies walk, fly, pursue mates, and search for food in diverse environmental and behavioral conditions, using only limited visual circuitry. How does this small, hardwired ensemble of neurons interpret such a wealth of constantly fluctuating visual information? A rich set of studies has demonstrated that sensory neurons encode the physical world dynamically, changing their filtering properties to suit varying sensory statistics [1, 2, 3]. However, despite evidence that cellular inputs to *Drosophila* motion detectors are capable of changing their processing properties [4, 5, 6], the circuit is frequently modeled as relying on inputs with static filters [4, 7, 8, 9, 10]. In this work, we use high-temporal resolution whole-cell patch clamp electrophysiology and modeling to ask two sets of questions. First, to what extent do the processing properties of inputs to *Drosophila* motion detectors change with varying visual statistics and behavioral states? In other words, do specific stimuli or states elicit more complex filtering waveforms? And second, given a dynamic input parameter space, can we define the fundamental computational motif underlying *Drosophila* direction selectivity?

In the *Drosophila* visual system, T4 and T5 are the first direction selective neurons (Figure 1A). They feed into multiple downstream pathways that control critical behaviors that depend on motion detection, such as course control [11], walking speed [12], and escape from looming stimuli [13]. T4 responds to ON motion while T5 is specific to OFF motion, with individual neurons of both types preferentially responding to local motion in one of four directions [14, 15, 7, 8]. We focused on the OFF pathway, where T5 receives columnar input (i.e. corresponding to one pixel in the field of view of the animal) from medulla cells Tm1, Tm2 and Tm4 in one column and from slower Tm9 cells in an offset, neighboring column (Figure 1A) [16, 17, 4, 18]. The connections between non-direction selective medulla cells and direction selective T5 cells constitute the locus of OFF motion computation [16, 17, 4, 18].

**Figure 1:**
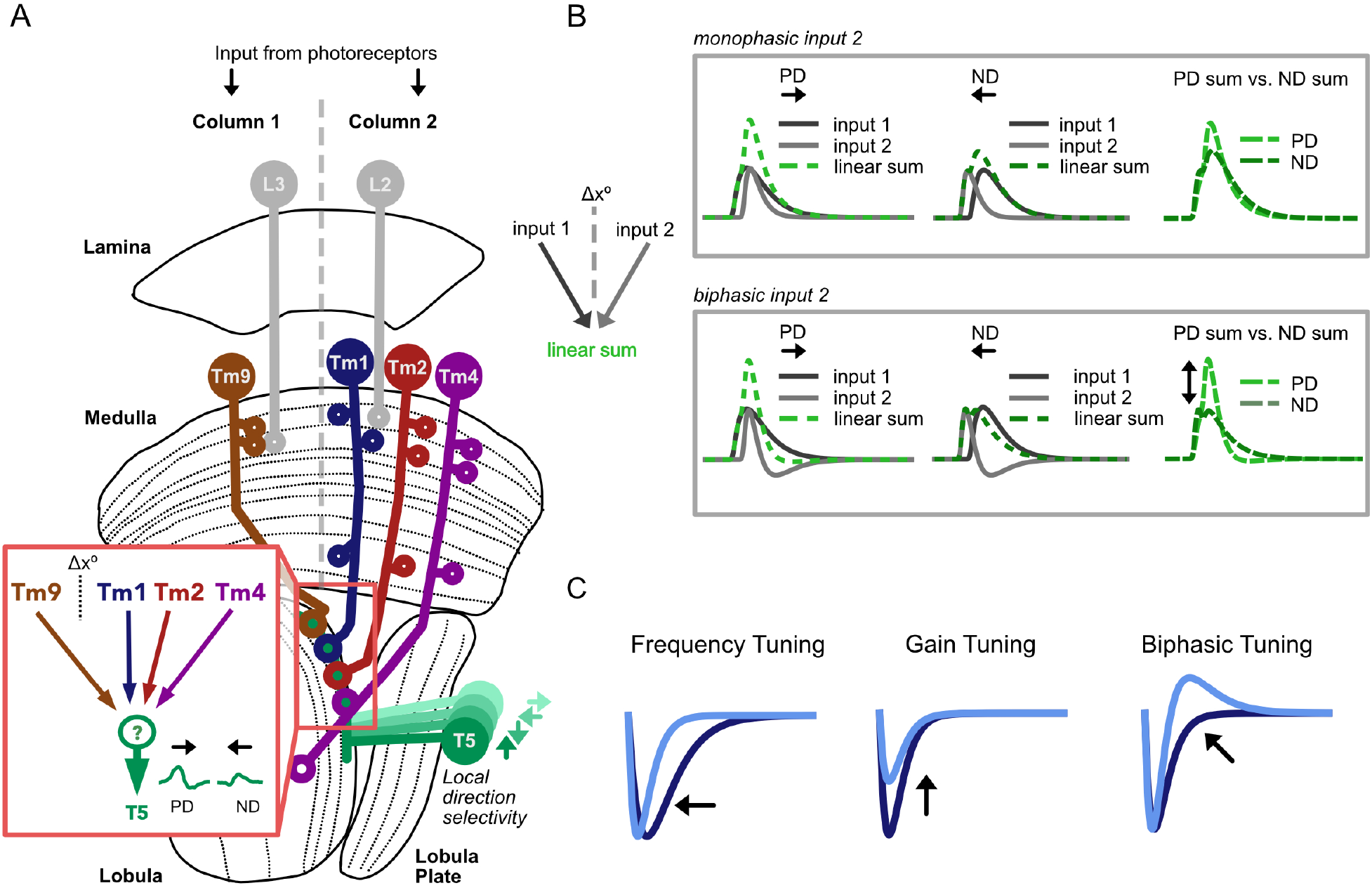
Motion detection in the *Drosophila* OFF Pathway. **A**. The connectome of the *Drosophila* OFF motion pathway is well characterized. *Inset:* T5s receive columnar input from Tm1, Tm2, Tm4 and Tm9. Tm1, Tm2, Tm4 (postsynaptic to lamina monopolar cell L2), and look at the same point in space. They are spatially offset (Δ*x*°) from Tm9 (postsynaptic to L3). Voltage responses in T5 are direction selective, depolarizing more strongly to motion in the preferred direction (PD) than to motion in the opposite, mnull direction (ND). The mechanisms underlying the emergence of these signals in T5 are debated. **B**. Filter shape can have a strong effect on the output of a motion detector. The linear combination of two spatially offset excitatory inputs, one of which is biphasic (*bottom*), can effectively suppress ND responses, generating an output that is more direction selective than the sum of two monophasic inputs (*top*). **C**. The temporal processing properties of sensory neuron filters have been shown to be stimulus and/or state dependent, varying in frequency, gain, and biphasic tuning.

Current models of direction selectivity at the level of T5 rely on a direct source of columnar inhibition, which is not supported by electron microscopy data [19, 18]. Furthermore, recent studies disagree on the fundamental computation underlying direction selectivity in T5, and argue for either a linear [20] or a nonlinear mechanism [4, 8]. However, these studies use different visual stimuli to probe the response properties of T5. We wondered whether the differences in stimuli between these studies could explain their disparate conclusions. In particular, since the temporal filtering properties of inputs to T5 govern the specificity and tuning of T5 output, differences in their shapes, which could be induced by different experimental paradigms, could drastically alter T5 responses. Most models of T5 do not measure input responses properties, and instead use simple monophasic (low-pass) temporal filters as inputs. However, linearly combining two spatially separated inputs, when one is biphasic (band-pass), can enhance direction selective responses (Figure 1B). It is therefore plausible that more complex spatiotemporal receptive fields in inputs can account for T5 direction selective responses, even in the absence of direct inhibition. Here we test this hypothesis by measuring stimulus- and state-dependent responses of the four main T5 columnar inputs.

Previous work has demonstrated changes in frequency tuning [4] and contrast gain adaptation [5, 6] at the level of T5 inputs. In order to look for signatures of changes to band-pass or “biphasic” tuning (Figure 1C), we used whole-cell patch clamp electrophysiology to record Tm1, Tm2, Tm4 and Tm9 responses to visual stimuli with a range of statistical properties. We also asked how stimulus-dependent properties might change in the presence of the neuro-modulator octopamine (OA), which is known to affect *Drosophila* motion circuits [21, 22, 4]. We found that the temporal properties of columnar inputs to T5 display stimulus- and state-dependent changes in filtering waveforms, including instances of strong biphasic responses. We used these stimulus- and state-dependent properties to characterize a “parameter space” for the filtering properties of the *Drosophila* motion vision circuit, through which these properties adjust dynamically to process changing visual statistics. We then linked the stimulus- and state-dependent responses of these inputs to previously measured T5 responses using a framework based on the *Drosophila* optic lobe connectome. Our model, based on a summation of the measured responses of excitatory OFF pathway medulla cell inputs, effectively recapitulates previously reported T5 responses when the model is adjusted to account for visual stimuli statistics. As such, our functionally and anatomically constrained model explains direction selectivity in T5 without the need for direct columnar inhibition. These results highlight the complex nature of this simple circuit, which endows an anatomically restricted set of neurons with the ability to encode a large space of stimuli, and shows that the question of mechanisms underlying direction selectivity is intimately intertwined with that of stimulus and state dependence.

## Results

### Filtering properties of columnar T5 inputs in response to a white noise stimulus

Linear-nonlinear (LN) models are widely employed to characterize spatiotemporal processing properties of individual cells in sensory systems [23, 24, 15, 4]. Following this approach, we first aimed to extract the linear spatiotemporal filters and associated nonlinearities that best describe the responses of Tm1, Tm2, Tm4 and Tm9 to a white noise stimulus, consisting of 5° horizontal bars (Figure 2 and Figure S1). From cellular responses to this stimulus, we extracted linear spatiotemporal receptive fields via reverse correlation [23, 25, 24, 4] and separated them into spatial and temporal components (Figure 2A-C). Our white noise results are in overall agreement with previous studies [24, 26, 17, 15, 4].

**Figure 2:**
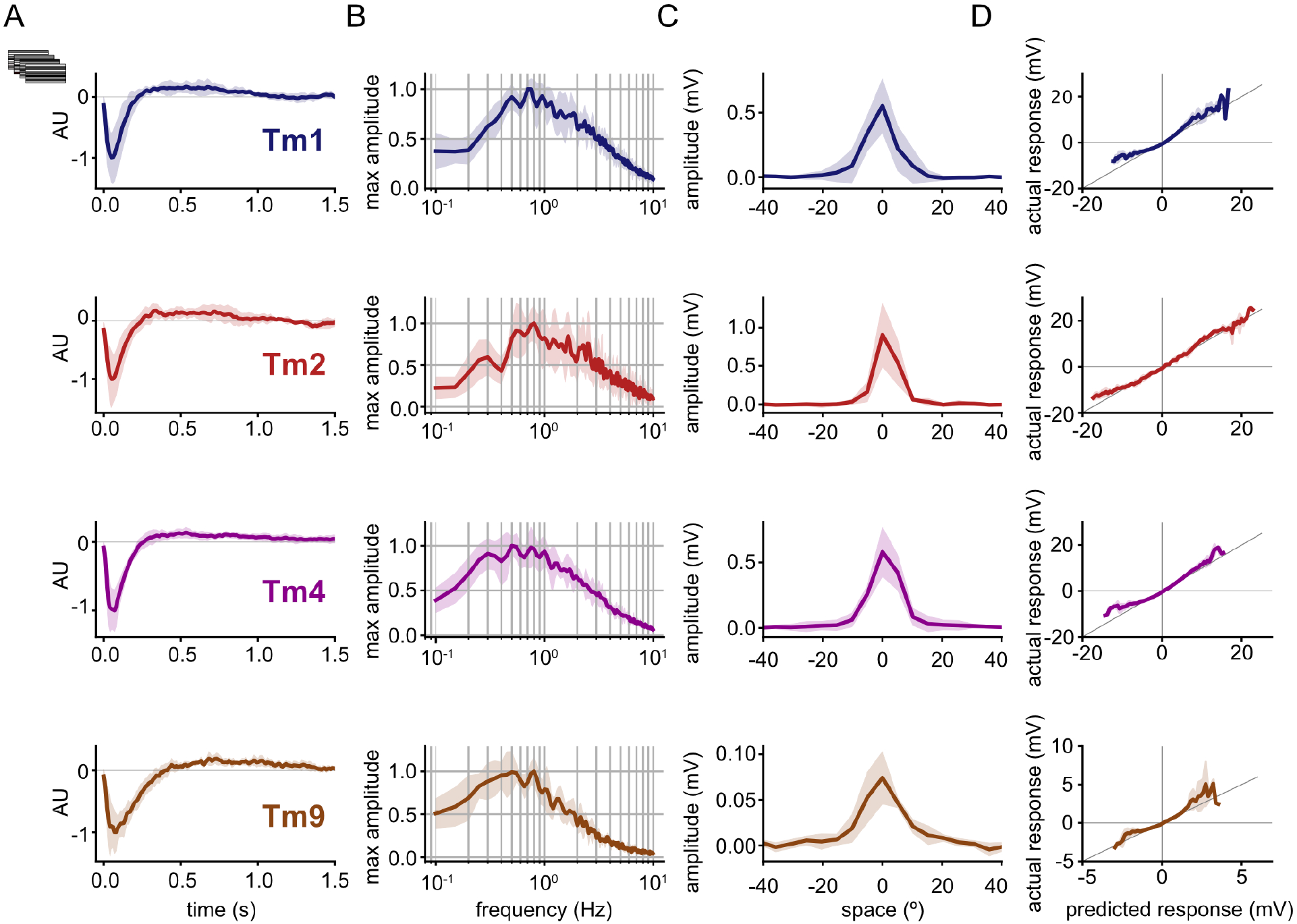
Tm1, Tm2, Tm4 and Tm9 exhibit spatial and temporal tuning with white noise stimuli. **A**. Normalized mean temporal filters for Tm1 (n=7), Tm2 (n=4), Tm4 (n=6), and Tm9 (n=6) extracted via white noise analysis. Filters show slight biphasic profiles. Shaded area represents standard deviation. **B**. Normalized mean frequency tuning of temporal filters when linearly convolved with sine waves of increasing temporal frequency. While all four Tm neurons are band-pass, Tm9 shows slower tuning than Tm1, Tm2 and Tm4. **C**. Mean spatial receptive fields extracted from spatiotemporal filters. **D**. Static nonlinearities for each cell type are linear for small voltage changes, but rectified at the upper and lower boundaries of their dynamic range.

As expected, the linear temporal filters of all columnar OFF-pathway inputs to T5 consist primarily of a negative lobe, indicating a sign-inversion between contrast polarity and cellular response (Figure 2A). The peak response times of our four filters are shorter than those reported by calcium imaging studies [4], and fall within a similar range to those found in previous studies of Tm1 and Tm2 using the higher temporal resolution techniques of either electrophysiology or voltage imaging [24, 27]. In the frequency domain, Tm1, Tm2, and Tm4 exhibit clear band-pass filtering properties, as previously noted [17, 4]. These band-pass properties correspond to the slight biphasic character of their linear temporal filters, which have shallow second positive lobes. In contrast to results obtained with calcium imaging, which determined that Tm9 is low-pass [26, 17, 4], we find that Tm9 also exhibits band-pass filtering properties (Figure 2B), albeit weaker than the other columnar inputs. Recording responses to long, 10 s flashes of light confirmed that these neurons are indeed band-pass as their responses return to baseline during the course of the stimulation (Figure S2).

We find that Tm1, Tm2, and Tm4 have narrow spatial receptive fields comprising approximately 10.8°, 8.2°, and 11.3° full width at half maximum (FWHM) when fit with Gaussians (Figure 2C, see Methods) [17, 4]. Tm9 has a slightly wider receptive field of approximately 15.3° FWHM [17, 4]. We found an additional subset of Tm9 cells that respond across a wide region of approximately 69.7° FWHM (Figure S3), as previously reported [26]. Tm9 responses fall naturally into two distinct populations based on their spatial receptive fields; however, with regards to their temporal properties, the two types of Tm9 responses are not distinct from each other (Figure S3). We therefore based our characterization of Tm9’s spatial properties on the population with narrower receptive fields, as these more closely matched the EM receptive field prediction from Shinomiya et al. [18]. Across cell types, we did not find center-surround structure in the spatial receptive fields extracted from our white noise stimulus (Figure 2C).

The static nonlinearities extracted from this dataset show that all four columnar T5 inputs respond linearly for small deflections in their membrane potential, but nonlinearly at the upper and lower boundaries of their dynamic ranges, with greater-than-linear depolarization amplitudes, and less-than-linear hyperpolarization amplitudes (Figure 2D). For stimuli that cause small deflections, the static nonlinearity only slightly improves fits (Figure S1). The contribution of static nonlinearity is more prominent with stimuli that cause large deflections such as high contrast flashes, where the negative components of the responses have lower amplitudes than the positive components (Figure S2). While this partial rectification is in line with previous studies for Tm1 and Tm2 [24, 27], calcium imaging studies have reported complete rectification for Tm1, Tm2 and Tm4 and a more linear response in Tm9 [17], in contrast to our findings.

These temporal and spatial filters, along with their associated static nonlinearities, offer a description of how Tm inputs to T5 process a white noise stimulus. But are they able to predict the responses of these neurons to stimuli with different visual statistics? To answer this question, we next recorded the responses of Tm1, Tm2, Tm4 and Tm9 to other types of visual stimuli that varied in time and contrast.

### Temporal processing of columnar inputs to T5 is stimulus dependent

We first recorded the response of the four columnar inputs to T5 to short full field brightness decrements of varying durations from a mean of grey. This type of stimulus has been previously used in the *Drosophila* motion detection circuit to characterize the response properties of Tm1 and Tm2 [27], as well as circuit output at the level of T5 [8]. Our aim was to compare these “flash” responses to predictions made from white noise filters. We measured the responses of Tm1, Tm2, Tm4, and Tm9 to high contrast flashes of 20, 40, 80, and 160 ms, and found that these did not match the output of our LN spatiotemporal white noise filters convolved with same stimuli (see Methods). These discrepancies appeared in both the shape and amplitude of the responses (Figure 3A). More specifically, we found Tm1 and Tm4 flash responses to be more biphasic than the corresponding white noise filter predictions across the four flash durations. Tm2 flash responses are more similar to the white noise prediction but also display a more biphasic response for 40 ms flashes. Furthermore, we find that the amplitude of the negative lobe of flash responses remains constant across stimulus durations for Tm1, Tm2, and Tm4. Shape-wise, Tm9 responses do not change significantly. Additionally, the gain of all Tm cell flash responses increases with flash duration. For Tm1, Tm2 and Tm4, white noise filter predictions of 20 and 40 ms flashes underestimate actual amplitudes of responses to flash stimuli, highlighting nonlinearities in gain at these shorter time scales. These comparisons indicate that temporal filters extracted in response to white noise are poor predictors of Tm1, Tm2, Tm4, and Tm9 responses to high-contrast flashes. In particular, the temporal responses of Tm1 and Tm4 are more biphasic under the latter stimulus conditions.

**Figure 3:**
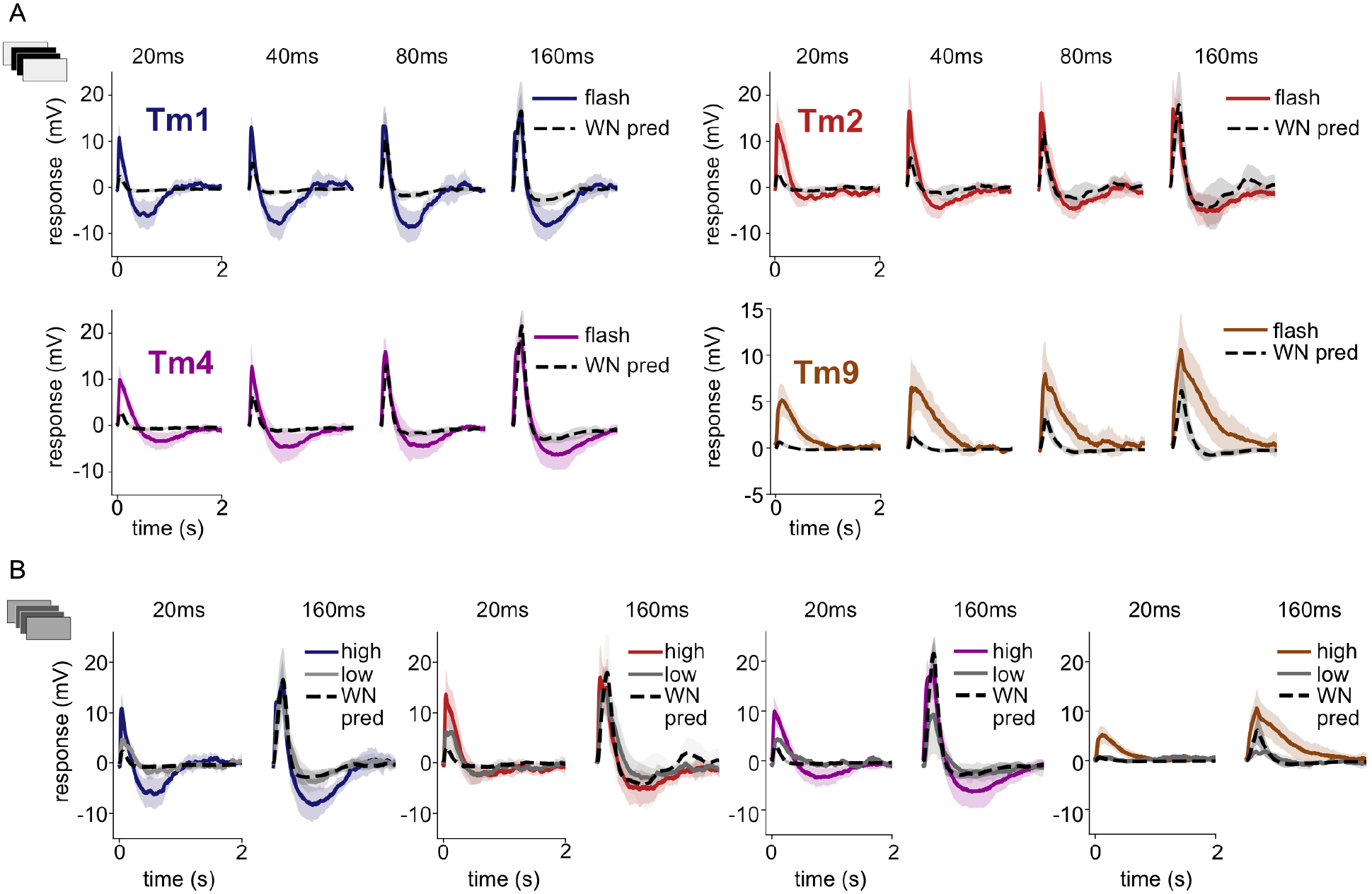
Tm1, Tm2, Tm4 and Tm9 responses are stimulus dependent. **A**. Mean Tm responses to 20, 40, 80 and 160 ms high contrast flashes (colored lines) are biphasic. Mean white noise filter predictions for the same 20, 40, 80 and 160 flashes (black, dashed lines) capture depolarization dynamics, but do not predict hyperpolarization for all flash durations, and under-predict depolarization amplitude for 20 and 40 ms flashes. Tm1 (n=4-5), Tm2 (n=5), Tm4 (n=5-6), and Tm9 (n=4-6). Tm9 white noise predictions fail to capture both depolarization amplitude and slower dynamics of Tm9 responses. Shaded area represents standard deviation. **B**. Mean Tm responses to 20 ms and 160 ms low contrast (grey) and high contrast flashes (color, same data as in B). Low contrast flashes do not elicit biphasic responses and are closer in shape to white noise predictions from A. Tm1 (n=4-5), Tm2 (n=4-5), Tm4 (n=1-2), and Tm9 (n=1-2).

We then sought to understand which statistical parameters of the stimulus drive the changes in response shape. The white noise stimulus, which changes contrast every 50 ms, falls within the same range of timescales as the flashes. However, a key difference between the white noise and flash stimuli is the magnitude of the contrast change. While the white noise stimulus consists of a truncated Gaussian distribution around the mean luminance of the projector (see Methods), the flash is a full contrast decrement from the same mean baseline luminance. Thus, a given contrast change for the white noise stimulus is, on average, smaller than that of the flash stimulus. To investigate the degree to which contrast could account for the change in biphasic properties of medulla cells, we measured responses of Tm neurons to flashes of lower contrast, starting at the same mean luminance level. We found that Tm1, Tm2, and Tm4 responses to flashes in low-contrast regimes lost their biphasic character, and more closely matched the white-noise filter predictions, both in terms of amplitude and waveform (Figure 3B). In the case of Tm9, which is only minimally biphasic to white noise, the response shape did not change significantly at different contrasts (Figure 3B, *right*). In general, high contrast flash responses are more biphasic than white noise responses, while low contrast flash responses are more comparable to white noise responses. We therefore find that the extent of contrast change of a flash stimulus is one factor in evoking the biphasic character of columnar T5 input responses.

We then asked if the results found using flash responses translate directly to a noise stimulus - i.e. does changing the contrast of a noise stimulus also affect the shape of extracted temporal filters? To answer this question, we recorded the responses of Tm1 to both high and low contrast ternary noise stimuli (Figure S4), consisting of random transitions between the mean luminance of the projector and fixed contrast increments/decrements of either high or low contrast, with the same temporal properties as the white noise. We find that Tm1 filters extracted from low and high contrast ternary noise have similar shapes to each other, as well as to the Tm1 filter extracted from white noise, presenting only as moderately biphasic. While we did not see a change in the shapes of filters, we did find that the amplitude of the temporal filter, or the gain, does increase with decreasing contrast, corresponding to an amplification of smaller signals that allows the cell to produce the same amplitude responses in different contrast regimes. Similarly, filters extracted from Tm2 and Tm4 responses to the high contrast ternary stimulus did not differ significantly in shape from filters extracted from the white noise stimulus (which was lower contrast), but had lower gains. While this finding is similar to that of Drews et al. [6] and differs from Matulis et al. [5], it confirms that a noise stimulus can indeed elicit changes in processing properties (i.e. gain) when its contrast statistics are modulated. However, when these results are considered in combination with responses to low contrast flash stimuli, which operate within similar contrast regimes, it becomes apparent that the shape of a temporal filter is stimulus dependent in a manner that is not only linked to contrast (see Discussion).

These results show that the temporal properties of columnar inputs to T5 are stimulus dependent and cannot be extracted using a single stimulus type or be fully described by a single temporal filter. Specifically, our data show that the shape of Tm1, Tm2, Tm4 and Tm9 responses are dependent on the statistics of the stimuli presented. In particular, specific stimuli can elicit strong biphasic temporal responses in Tm1, Tm2 and Tm4.

### Temporal processing of columnar inputs to T5 is state dependent

Exploring stimulus-dependent changes in processing revealed that inputs to T5 change their filtering properties in response to different visual stimuli. Are the changes in stimuli able to capture the full range of response properties that these cells can generate? To answer this question, we next investigated state-dependent changes, which have also been found to dramatically affect the encoding properties of sensory neurons through the action of small molecule neuromodulators [28, 29, 30]. The neuromodulator octopamine (OA), released during locomotion, has been found to change the gain and frequency tuning of T4 and T5 cells as well as downstream lobula plate tangential cells (LPTCs) [21, 22, 4]. These changes are likely inherited in part through the modulation of columnar transmedullary inputs to these circuits. Indeed, Arenz et al. [4] report an acceleration of the responses of inputs to T4 and T5 with application of CDM, an OA agonist. We therefore investigated how these state-dependent changes in processing properties compare with the stimulus-dependent changes that we observed, and whether any relationship could be drawn between the two.

We conducted recordings in the presence of the neuromodulator OA, and found that the application of 10 *µ*M OA induced a strongly biphasic white noise extracted temporal filter in Tm1, Tm2 and Tm4, with a sharp, positive second lobe emerging (Figure 4A). Corresponding to the emergent biphasic character induced in each cells’ linear temporal filter, responses are more band-pass with OA. In addition, OA induces faster temporal filter peaks for Tm1, Tm2 and Tm4. This latter effect, also apparent in Arenz et al. [4], manifests in the frequency domain as a shift toward higher frequencies (Figure 4B). The frequency tuning of Tm9 does not change significantly in the presence of OA (Figure 4A-B). As previously noted by Arenz et al. [4], the application of OA did not exert any significant effect on the spatial receptive fields of any columnar T5 input (Figure 4C). Additionally, while the application of OA reduces the gain of these cells, it does not appear to significantly change their static nonlinearities (Figure 4D). This reduced dynamic range in the presence of OA likely prevents them from reaching response amplitudes at which nonlinear processing effects are seen (Figure 4D).

**Figure 4:**
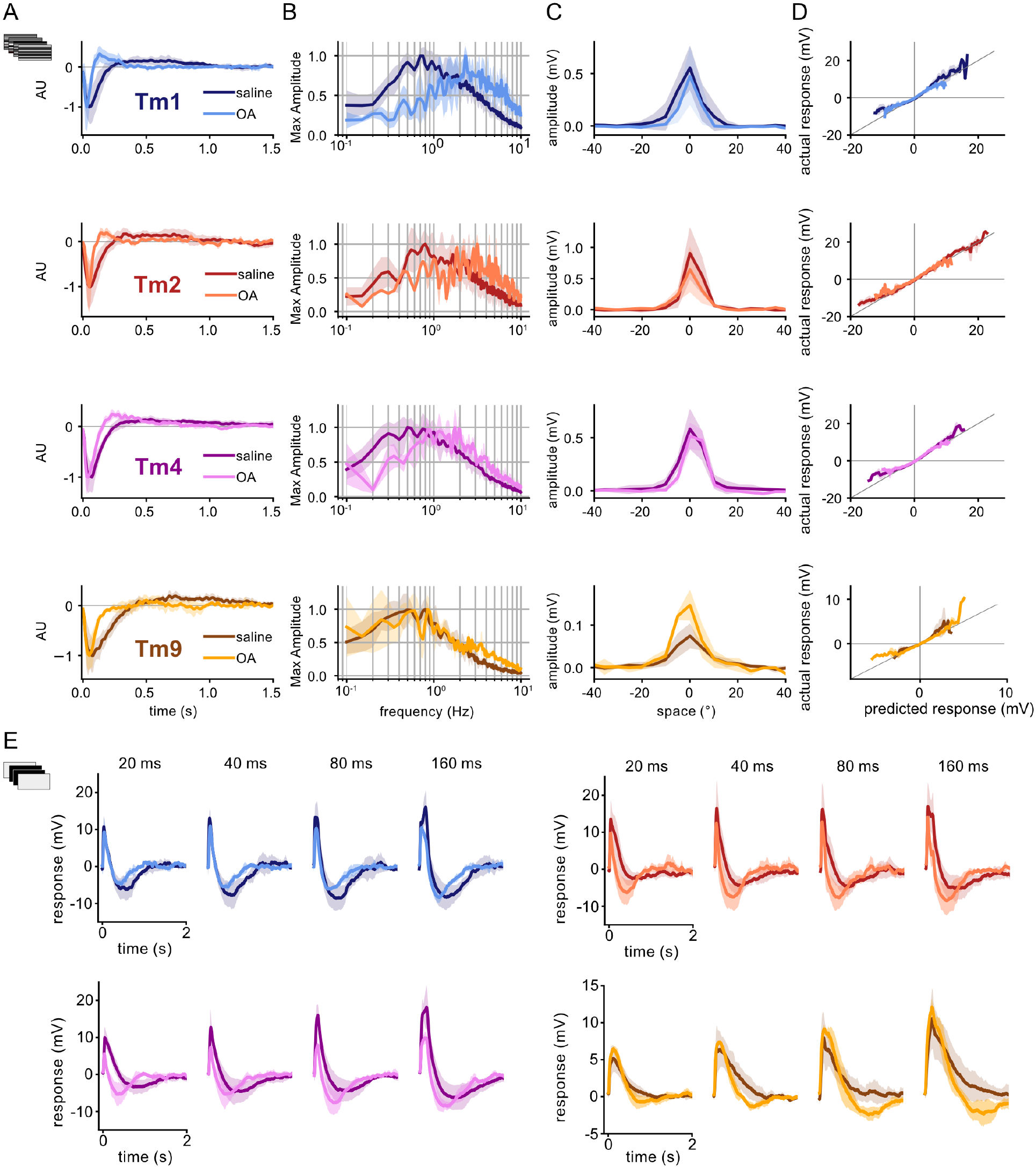
Tm1, Tm2, Tm4 and Tm9 exhibit stimulus and state dependence in presence of neuromodulator octopamine (OA) Responses in presence of OA (lighter colors) are overlaid with data from Figures 2-3 (darker colors) **A**. Temporal filters extracted from white noise stimulus show distinct changes in shape, with a faster and narrower first lobe and the emergence of a sharp second lobe in the case of Tm1, Tm2 and Tm4 in OA. **B**. In frequency space, Tm1, Tm2 and Tm4 filters in the presence of OA show distinct shifts in tuning to higher frequencies. Tm9 tuning does not change. **C**. There are no significant changes in spatial filters -/+ OA. **D**. There are no significant changes in static nonlinearities between conditions. **E**. Tm1 (n=3-4), Tm2 (n=3), Tm4 (n=3-4), and Tm9 (n=2-3) responses to flashes (20, 40, 80, 160 ms) are faster and, in most instances, more biphasic in OA.

We next asked how the effect of OA interacts with the stimulus-dependent changes in processing properties we had previously observed in columnar T5 inputs, and assessed the effect of OA on responses to flash stimuli. In addition to having faster kinetics, flash responses in the presence of OA were more biphasic for all four cells as compared to saline conditions (Figure 4E). The corresponding white noise predictions made using the temporal filters extracted in OA are poor predictors of these flash responses (Figure S5A). Similar to saline conditions, low contrast flashes produced less biphasic responses than high contrast (Figure S5B); however, low and high contrast flashes in OA are both more biphasic than those measured in saline. This reveals a nuanced relationship between state- and stimulus-dependent changes to processing properties, where state and stimulus can elicit similar shifts both independently or in tandem with one another.

Neuromodulation therefore has a strong effect on the shape of the temporal responses of columnar inputs to T5. In OA, the temporal properties of Tm1, Tm2 and Tm4 not only become faster, shifting their tuning towards higher temporal frequencies, but also display changes in the waveforms of their temporal filters, which become more biphasic.

### T5 columnar temporal responses move across a continuous stimulus- and state-dependent parameter space

The similarities between shape changes in the temporal responses of T5 columnar inputs to either high contrast flashes or to responses measured in the presence of OA hinted at a continuum of responses rather than discrete differences (Figure 5A-D). To describe this stimulus- and state-dependent “parameter space” of responses for each of the inputs to T5, we compared all stimulus and state conditions on a similar temporal timescale, using parameterized responses (Figure S6, see Methods).

**Figure 5:**
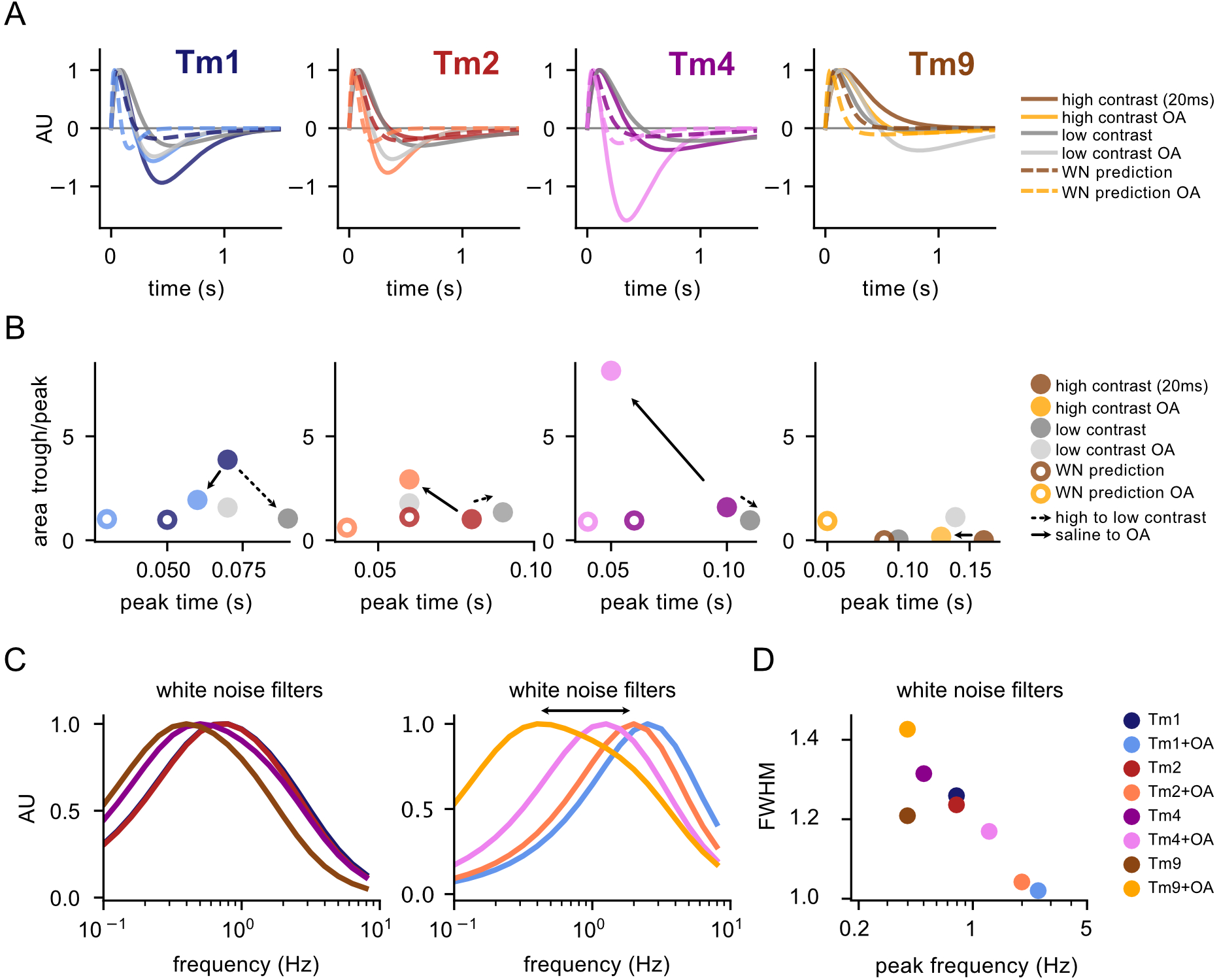
Tm1, Tm2, Tm4 and Tm9 temporal responses lie within a parameter space. **A**. Comparison of parameterized Tm1, Tm2, Tm4 and Tm9 responses to 20 ms flashes across conditions, including high contrast and high contrast with OA (solid colored lines), low contrast and low contrast with OA (grey lines), and both baseline and OA LN white noise filter predictions for 20 ms stimuli (dashed lines). **B**. The parameterized responses from A are plotted as a function of peak time (x-axis) vs. the ratio of the area of the peak lobe with respect to the trough lobe (y-axis). **C**. Frequency tuning of parameterized baseline (*left*) and OA temporal filters (*right*). Filters in OA become more band-pass and shift their peaks to higher frequencies (black arrow). **D**. Full width half max (FWHM) as a function of temporal frequency of filters in C.

We focused on responses to a 20 ms flash stimulus, either measured directly or predicted from white noise filters across all Tm cells, in the absence and the presence of OA. Normalized and plotted together (Figure 5A), it is clear that Tm1, Tm2 and Tm4 exhibit a wide range of responses, while Tm9 shows somewhat fewer changes across stimuli and state. To better visualize how different conditions affect these responses, we plotted the ratio of the area of the trough by the area of the peak as a function of peak time, roughly representing the extent of a filter’s biphasic character as a function of speed of response (Figure 5B). The 2D space occupied by the Tm neurons within this plot illustrates the span of the diversity of responses within cell types and reveals global trends: responses move from being less to more biphasic between noise and flash stimuli, and shift toward being faster and more biphasic in the presence of OA. In the case of white noise filters, the effect of OA is particularly clear in the frequency domain (Figure 5C). In the case of Tm1, Tm2, and Tm4, OA shifts peak responses towards higher frequencies so that their frequency tuning curves are spread further from each other (and thus across a broader spectrum of frequencies) compared to saline conditions. Additionally, OA decreases the tuning curve FWHM values, thereby making the OA filters more band-pass (Figure 5D).

High temporal resolution electrophysiological recordings of Tm1, Tm2, Tm4 and Tm9 under different stimuli and neuromodulatory conditions reveal a highly flexible circuit with the ability to display changes in temporal filter shape. We next investigated the computational consequences of these stimulus- and state-dependent properties.

### A sum of columnar inputs predicts T5 flash responses

Recently, Gruntman et al. [8] obtained whole-cell patch clamp recordings of T5 responses to stationary high contrast flashing bars. The authors found T5 to display asymmetric hyperpolarizing responses. For any particular T5 cell, flashing bars on the side of the spatial receptive field corresponding to the leading edge of the cell’s preferred direction of motion elicited only a depolarizing response. Bars on the opposite side of the receptive field, however, caused a depolarization followed by a hyperpolarization. To explain these results, the authors propose a direction selective model in which the functional properties of T5 can be explained by a combination of direct columnar excitation and inhibition. Since no such columnar inhibitory input has been found by connectome studies [18], we instead hypothesized that the strongly biphasic nature of the temporal responses of Tm1, Tm2 and Tm4 to flashes could explain T5 responses without the need for a direct inhibitory input. Because Tm1, Tm2, and Tm4 have similar processing properties (Figure 2D-G) and look at the same point in space [18], we asked whether a single biphasic excitatory columnar input combined with Tm9 via linear regression could capture the dynamics of the T5 response, and the asymmetric hyperpolarization in particular.

We used our measured responses of Tm1 and Tm9 to predict T5 responses to stationary high contrast flashing bars without additional manipulation. To compare our data with existing T5 data, we first convolved the white-noise extracted linear temporal filter of each cell type with a 1D stimulus of length 20, 40, 80 and 160 ms [7, 8]. Using linear regression with positivity constraints, we fit these predicted responses to T5 flash responses collected by Gruntman *et al*.[8]. As expected from their shape, we found that the white noise filter predictions were able to capture the depolarizing responses, but failed to capture the asymmetric hyperpolarization (Figure 6A, *top*).

**Figure 6:**
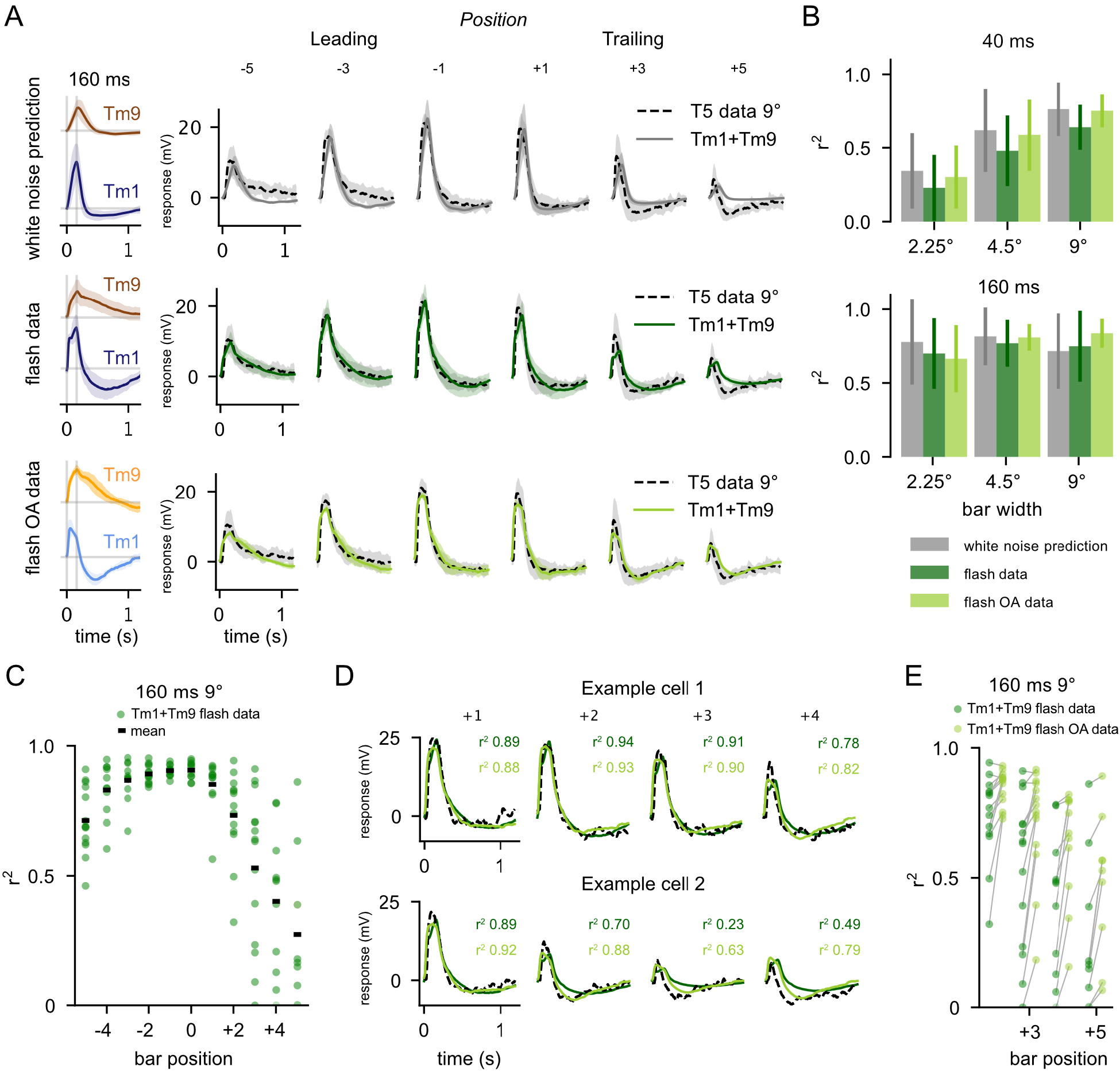
The sum of columnar inputs predicts T5 flash responses. **A**. *Top:* White noise extracted filters are convolved with 160 ms stimulus and then fit with linear regression to T5 electrophysiological recordings from Gruntman et al. [6] for the 160 ms, 9° condition, at various positions in the receptive field of T5 (data dashed line, fit solid grey line). T5 average traces shown for bar position from “Leading” edge (*—* 5, *−* 3, *−*1) and “Trailing” edge (+1, +3, +5). *Middle:* Average Tm1 and Tm9 responses to 160 ms flashes are fit via linear regression to each T5 recording from Gruntman et al. [6] for the 160 ms, 9° condition (data dashed line, fit solid dark green line) *Bottom:* Same as *Middle* using Tm1 and Tm9 160 ms flashes in the presence of OA (data dashed line, fit solid line) Linear regression using flash responses and flash responses recorded in OA provides a good fit to T5 data. This is especially evident in the trailing edge (bar positions +3 and +5). **B**. Aggregate *r*^2^ values (square of sample correlation coefficient, see Methods) across bar positions for linear regression fits of Tm1+Tm9 to Gruntman et al. [8] recordings of T5 (conditions: 40 and 160 ms presentations of 2.25°, 4.5°, and 9° bars). Error bars depict standard deviation **C**. Distribution of *r*^2^ values across bar positions for fits to individual T5 responses to 160 ms, 9° bars. **D**. Example traces of fits to two single cells from C (T5 data, black dashed line; fits using saline flashes, dark green; fits using OA flashes, light green). **E**. Using the highly biphasic Tm1/Tm9 flashes recorded in OA improves the *r*^2^ of fits on the trailing edge of the T5 receptive field, where asymmetric hyperpolarization is most evident.

We next asked if our flash responses, which were obtained from an experimental paradigm more similar to the single-position bar flashes of Gruntman *et al*., could predict the full response properties of T5 more accurately than our white noise filters. Using linear regression with positivity constraints, we found that a weighted sum of Tm1 and Tm9 responses derived from flash stimuli do better at reproducing measured T5 responses to single-position bar flashes (Figure 6A, *middle*), but still fall short of capturing both the extent and the kinetics of T5’s asymmetric hyperpolarization at the trailing edge of the T5 receptive field. Since Tm1 flash responses obtained in OA conditions have faster kinetics and larger second lobes, we also ran the linear regression using flash responses of Tm1 and Tm9 obtained in the presence of OA, despite the apparent mismatch in recording conditions. In this case, the linear regression provides a near perfect fit with T5 data (Figure 6A, *bottom*).

It was puzzling that the flash responses recorded in OA provided such a good fit in the linear regression, as Gruntman et al. [8] acquired these data in regular saline and not in OA-supplemented saline. It is, however, conceivable that endogenous state modulation occurred during T5 recordings. This was hinted at by the apparent variability in the amplitude and kinetics of the asymmetric hyperpolarization in T5 responses across different cells [8]. To investigate this, we performed linear regression on individual T5 cells, instead of the average of all recordings, using flash responses recorded in either saline or saline with OA. For a subset of T5 cells, which displayed a slower and less salient hyperpolarization, the saline linear regression provided a good fit (Figure 6C and D *top*). For a different set of T5 cells, the OA linear regression provided a better fit (Figure 6D *bottom*). This indicates that the diversity of responses in the T5 data largely accounts for the distribution of our *r*^2^ values (Figure 6C). In these cases, performing the linear regression using the OA flash responses often increased the *r*^2^ value substantially (Figure 6E). Although we performed this analysis using Tm1 and Tm9 to predict 9° 160 ms T5 flashes, these results stand across flash durations and widths (Figure S7A) as well as using other combinations of Tm inputs (Figure S7B).

In all conditions, the coefficients output by this linear regression show distinct separation between Tm1 and Tm9 (Figure S7C), similar to that seen in the electron-microscopy (EM) data. In addition, the weighted spatiotemporal receptive fields constructed by linearly combining Tm1 and Tm9 fits are tilted in space-time, indicating direction selectivity. The tilt in space-time is more prominent when these are constructed from flash responses, both in saline and OA, demonstrating the increased effectiveness of flash responses at capturing T5 direction selectivity. In agreement with this, the same linear regression fits predict the profile of T5 responses to moving bars from Gruntman et al. [8], as well as direction selectivity (Figure S7D, see Methods).

These results demonstrate that including a biphasic input to T5 can account for its spatially asymmetric hyperpolarization. This shows that stimulus- and state-dependent properties of inputs strongly affect output at the T5 level. Thus, for a model to accurately describe the direction selective responses of T5 to moving stimuli, it must consider both stimulus- and state-dependent processing properties of its inputs.

### A connectome-based model captures OFF pathway direction selectivity in the context of different stimuli and states

Motivated by the linear regression, we built a model of T5 direction selectivity that is faithful to the anatomy of the circuit and takes into account our experimental measurements of Tm response properties. We imposed the following overarching constraints: (1) T5 receives inputs from Tm9 in one ommatidial column, and Tm1, Tm2, and Tm4 from an adjacent column, (2) all four T5 inputs are excitatory (cholinergic), and (3) the response properties of the transmedullar inputs vary with stimulus or state, as we demonstrated. We captured the first constraint by separating the center of the receptive field of Tm9 by 5° from the rest of the Tm cells (Figure 7A). The second constraint was satisfied by requiring all cells to provide positive input to T5. Additionally, we used the relative synaptic counts of Tm1, Tm2, Tm4 and Tm9 from the connectome as synaptic weights to constrain the relative contribution of each cell type to T5 responses [18]. As for the third constraint, when constructing the four inputs to T5, we matched their response properties with the stimulus presented to our model, such as moving sine waves [20, 8] or high contrast moving bars [8].

**Figure 7:**
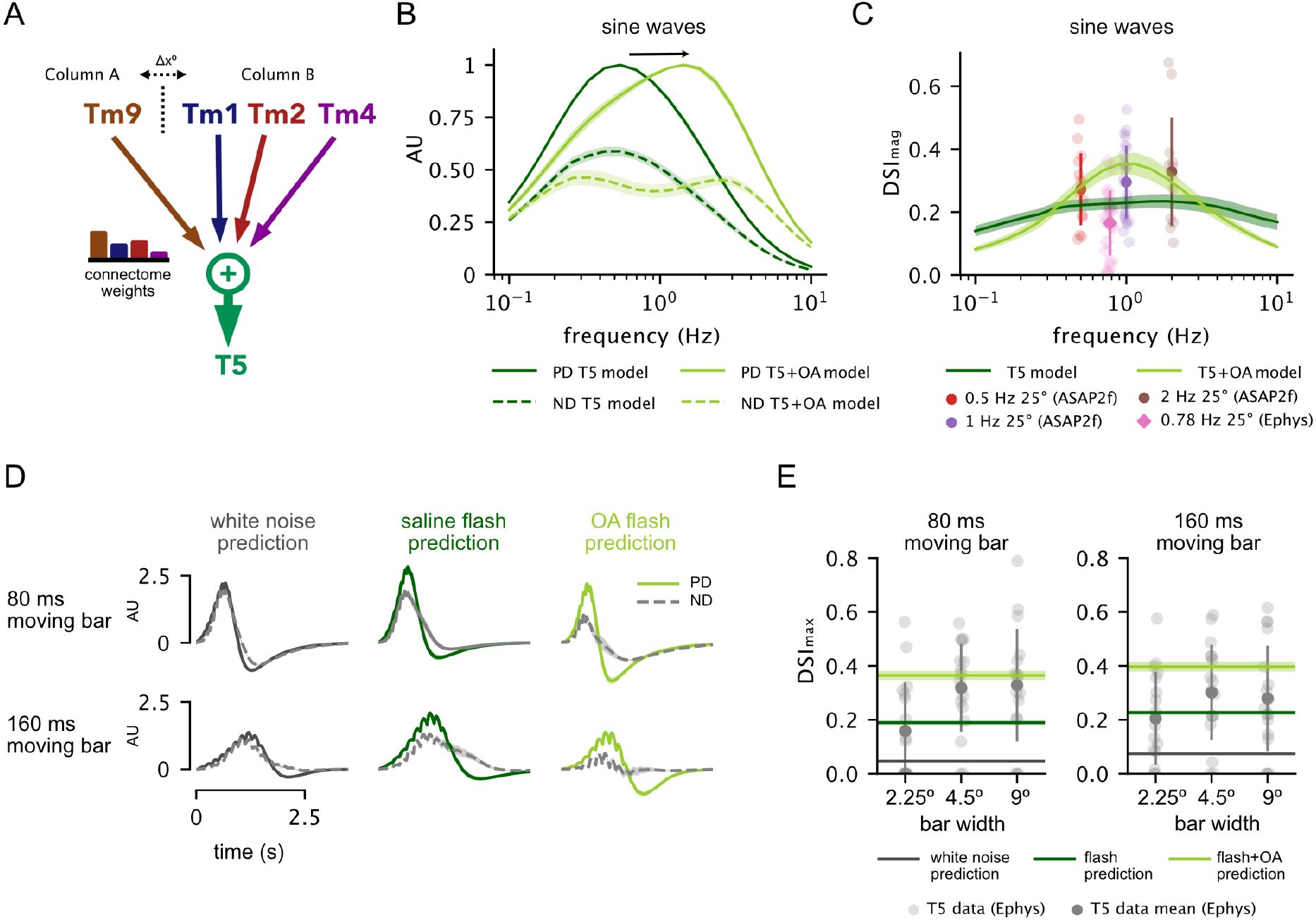
Low-Parameter, Connectome Based Model is Sufficient to Capture OFF Pathway Direction Selectivity in the context of different stimuli and states. **A**. Schematic of model framework constructed with Tm9 spatially offset from Tm1, Tm2 and Tm4 by Δ*x* = 5°. **B**. Preferred direction (PD) and null direction (ND) frequency tuning of model to sine waves using parameterized spatiotemporal filters extracted in saline alone vs. those extracted in the presence of OA. **C**. Direction selectivity index (DSI_mag_ = (|PD|*−* |ND|)*/*(|PD| + |ND|), see Methods) for model using saline-derived filters with n=20 samples of published EM weights from [16] across various frequencies (dark green green) compared to output using OA-derived filters (dark green). Experimental voltage-imaging (ASAP2f) T5 DSI data shown from Wienecke et al. [20] (circles), and T5 electrophysiology data from Gruntman et al. [8] (diamonds). **D**. Example PD and ND model output traces for an 80 ms and a 160 ms moving bar stimulus, with inputs based on white noise predictions (*left*, black), flash responses recorded in saline (*middle*, dark green) and OA (*right*, light green). **E**. Using flash response-based inputs, model DSI falls within the range of T5 electrophysiology data reported by Gruntman et al. [8] for moving bars. Direction selectivity index (DSI_max_ = (max(PD) *−*max(ND))*/*(max(PD) +max(ND), see Methods) increases when using OA-based flash responses due to their biphasic nature.

We first modeled T5 responses to sine waves. To describe the response of each T5 input to this stimulus, we used the temporal and spatial filters of Tm1, Tm2, Tm4, and Tm9 extracted from white noise analysis, as well as their associated static nonlinearities (see Methods). These filters accurately predicted measured responses of Tm cells to sine waves (Figure S8), making them appropriate descriptors of cellular responses in this particular stimulus regime. Output from this model in response to sine waves matched T5 data from previous studies, in that it predicted maximum preferred direction (PD) tuning just below 1 Hz (Figure 7B) [21, 4]. The direction selectivity index (DSI) for the output of the model also fell within the range of experimentally calculated DSIs from two recent studies: Wienecke et al. [20], using voltage-imaging, and Gruntman et al. [8], using electrophysiology (Figure 7C). We then asked how the enhanced biphasic character and shifted frequency tuning of filters extracted in the presence of OA affected model output. In this case, our model predicted a broadening and a shift in T5 PD frequency tuning toward faster frequencies (Figure 7B) that matched previous measurements of T5 [4] and LPTC [22] tuning in the presence of OA or the OA agonist chlordimeform (CDM). Furthermore, using OA-derived filters increased DSI (Figure 7C). These results using white noise filters show that combining Tm1, Tm2, Tm4 and Tm9 responses linearly with EM connectome weights is sufficient to achieve the direction selective response of T5 cells to sine waves across studies.

We next modeled T5 response to moving high contrast bars. The results of our linear regression analysis showed that strongly biphasic Tm responses best predicted T5 flashing bar responses. As expected, the characteristic white noise filters for Tm1, Tm2, Tm4 and Tm9 did not capture the DSI of T5 responses to moving bars (Figure 7D *left*, E). We therefore constructed a corollary model of T5 based on parameterized flash responses for Tm1, Tm2, Tm4 and Tm9 (see Methods). The increased biphasic nature of the flash responses allowed the model to achieve direction selectivity for moving bar stimuli in the range of T5 recorded electrophysiology data [8] (Figure 7 D *middle*, E). In this case, the negative lobe from strongly biphasic Tm inputs cancels out depolarizations in lieu of direct inhibition. Correspondingly, flash responses obtained in the presence of OA, which are more biphasic than those obtained in saline alone, increased the model’s DSI when used as inputs (Figure 7 E *right* and F).

These results demonstrate that the increased biphasic character of Tm cells, which occurs both as the result of changes to stimulus or the presence of a neuromodulator, can produce direction selectivity on par with that seen in T5 electrophysiology recordings.

Previous studies were unable to reconcile direction selectivity in T5 with the constraint of solely excitatory columnar inputs due to underdescribed processing properties of Tm cell inputs to T5, leading them to invoke an illusive source of direct inhibition. Our state- and stimulus-dependent measurements and modeling therefore reconcile anatomy and function in a canonical *Drosophila* motion circuit.

## Discussion

In this study, we demonstrated that the response properties of neurons in the *Drosophila* OFF motion pathway are shaped by both visual stimulus statistics and a behaviorally relevant neuromodulator, and that such flexibility clarifies the computation of direction selectivity. We found that Tm white noise-extracted filters can be poor predictors of Tm cell responses to stimuli with different visual statistics. Specifically, these filters fail to capture changes in the shape of the responses to high contrast flashes. We also demonstrated that similar changes to the filtering properties of T5’s columnar inputs arise due to the action of the neuromodulator octopamine. Incorporating these state- and stimulus-dependent properties into an anatomically constrained model of the motion circuit based on input summation yields a good prediction of T5 responses across stimulus regimes. Our results demonstrate that neurons in the *Drosophila* visual system operate within a stimulus- and state-dependent space of temporal filtering parameters, and are underdescribed by the filters commonly used in *Drosophila* motion circuit models.

### Stimulus- and state-dependent changes in filtering properties highlight circuit flexibility

Flexible processing of stimuli is a ubiquitous feature of sensory systems across species [1, 2, 3]. In blowflies, the temporal properties of lamina monopolar cells (LMCs), the main inputs to the transmedullary cells that we focused on in this study, display stimulus-dependent changes in shape. Both van Hateren [31] and Srinivasan et al. [32] have shown that LMC responses are more biphasic with increasing signal to noise ratio (SNR) of the stimulus. These studies provide a rationale for differences across conditions. A monophasic, or low-pass, filter acts as an integrator, extracting slow temporal components of a visual scene. This is useful when visual information is noisy (low SNR), because increases in the redundancy of information translate into increases in reliability. A biphasic, or band-pass, filter, however, is advantageous in high SNR conditions because it acts as a differentiator and efficiently conveys changes in the stimulus, thereby decreasing correlations and reducing redundancy [33].

When comparing responses across stimuli in different SNR regimes, our recordings of Tm1, Tm2 and Tm4 are compatible with these hypotheses. The temporal filters of these three neurons have less biphasic shapes in response to temporally unstructured stimuli that have the characteristics of noise, both white and ternary, which we consider to correspond to a low SNR regime. Responses to low contrast flashes, which can also be considered low SNR, are also close to monophasic and are well predicted by white noise filters. On the other hand, high contrast (high SNR) flashes produce strong biphasic responses. The properties of Tm1, Tm2 and Tm4 are therefore similar to and likely inherited from their LMC input (primarily L2, Figure 1A).

Interestingly, we find that the addition of OA produces a more biphasic character in the white noise-extracted temporal filtering properties of Tm1, Tm2, and Tm4, similar to the waveform changes seen in response to high contrast flashes. Following the logic discussed above, this change in filter shape would optimize information processing in high SNR regimes. More biphasic, differentiator-like responses may be beneficial for the rapidly changing visual scene when a fly is walking or flying. This raises the possibility that OA, by shifting processing properties towards those induced by high SNR stimuli, acts to prime T5 inputs to detect salient stimuli in the natural statistics of a moving or flying fly. Furthermore, columnar inputs to T5 express receptors for many neuromodulators other than OA [34], suggesting that state-dependent modulation of motion detection likely plays an even more heterogeneous role, with multiple neuromodulators acting in concert at any given time.

In addition to changes in filter shapes, we observed OA-dependent shifts in the kinetics of the temporal filters of Tm1, Tm2 and Tm4 towards faster speeds. Locomotion, through the release of OA, has previously been shown to broaden and shift the tuning of *Drosophila* motion detector outputs toward higher frequencies [21, 22]. This mechanism is thought to tune motion pathways to the increased frequencies of motion that flies experience as a result of self-motion during locomotion. This effect is thought to be, at least in part, through the modulation of T4/T5 inputs [4]. Our findings corroborate the hypothesis that octopaminergic modulation of frequency tuning in this circuit is inherited in part from upstream elements. In addition, our high temporal resolution data shows that Tm1, Tm2 and Tm4 have similar temporal response dynamics in saline, but acquire different kinetics in the presence of OA. This broadens the range of temporal frequencies collectively encoded by these three neurons (Figure 5A), an effect that we see in the output of our model. Thus, while Tm1, Tm2 and Tm4 might appear to have redundant roles, the differential effect of OA on these three T5 inputs highlights a functional relevance in the context of changing behavioral states. Finally, in contrast to Tm1, Tm2, and Tm4, we find the temporal filtering properties of Tm9 to be less affected by either stimulus statistics or by the presence of OA.

We focused here on changes in temporal dynamics; however, it is likely that additional processing properties of Tm neurons, such as in their spatial receptive fields, are sensitive to both stimulus and state. Integrating changes in these processing properties could hypothetically fine-tune the motion-selective outputs across conditions. In addition, we find two distinct classes of Tm9 cells with different sizes of receptive field, as has been previously reported [26]. Although larger spatial receptive fields may not contribute directly to direction selectivity, further characterization of this heterogeneity may provide insight into diverse T5 responses.

### Accounting for stimulus dependence clarifies circuit mechanisms

Although direction selectivity has been investigated since the 1950s, the mechanisms underlying motion detection in the invertebrate visual lobe and their cellular implementation are still being debated [35, 36]. For the OFF pathway that we have explored, one debate concerns the linearity of the summation of inputs to directionally selective T5 neurons. Wienecke et al. [20] argue that the response of T5 axonal terminals to stationary and moving sine waves suggests linear summation, whereas Gruntman et al. [8], who studied responses to flashed and moving bars, argue for nonlinear summation. Neither of the studies had access to the waveform of the actual inputs to T5 - the results we have presented. On the basis of this additional knowledge, our modeling work supports linear summation. In addition, although T5 responses show apparent suppression in some regions of the visual field, we find that this does not require an inhibitory input. Instead, the biphasic character of the Tm1, Tm2, and Tm4 responses can reproduce the data without direct inhibition. Furthermore, we found that the model could account for direction selectivity when not only the identity but also the strengths of its connections were determined directly from the connectome data [18].

It should be stressed that we are not proposing that inhibition plays no role in the directionally selective OFF pathway. For example, the wide-field inhibitory cell CT1 [37, 18] may provide wide-field gain normalization [38, 6]. Such normalization could enhance direction selectivity, but we argue that it is not necessary for producing it.

### Stimulus and state dependence in sensory processing of natural scenes

It is well established that white noise filter characterizations of cells in the mammalian retina and V1 are poor predictors of responses to natural scenes [39]. We expect that Tm white noise filters will similarly fail to capture Tm responses to natural scenes. However, it is likely that responses to natural scenes will occupy the “parameter space” defined by the diverse responses probed here and elsewhere [27, 35, 4, 6, 5]. Many approaches have been proposed for characterizing cell responses to natural stimuli in an interpretable manner [40, 39, 41]. Correctly incorporating the link between scene statistics, the location in parameter space, and the appropriate Tm filtering properties will be essential in accounting for direction selectivity in a natural setting.

## Acknowledgments

We thank A.J. Zimnik and S.L. Heath for comments on the manuscript. We thank C.F.R. Wienecke and T.R. Clandinin for sharing T5 sine wave data. J.R.K. acknowledges support from the James H. Gilliam Fellowships for Advanced Study program (HHMI) and NIH F31EY030319. J.P.P was supported by Neuronex NSF 1707398. M.P.C. was supported from NIH 5T32EY013933 and NIH R01EY029311. L.F.A was supported by NSF NeuroNex Award 1707398, the Gatsby Charitable Foundation GAT3708 and the Simons Collaboration for the Global Brain. R.B. was supported by NIH R01EY029311, the McKnight Foundation, the Grossman Charitable Trust, the Pew Charitable Trusts, and the Kavli Foundation.

## Methods

### Electrophysiology Preparation

In order to target specific medulla cell populations, we used the Gal4-UAS binary expression system in a w+ background to drive expression of a cytosolic variant of GFP in each individual fly line. Gal4 lines were as follows: R71G04-Gal4 (Tm1), otd-Gal4 (Tm2), R35H01 (Tm4), and R24C08-Gal4 (Tm9). Live flies were immobilized in a position that allowed them to see visual stimuli while providing physical access to one optic lobe, and electrophysiological recordings were performed as in Behnia et al. [24].

### Stimulus Presentation

We built visual stimuli using our own custom extension of the Allen Brain Institute’s retinotopic-mapping package [42]. Each stimulus was warped and projected onto a flat screen aligned with the left eye. To correctly warp the stimulus, we assumed the eye was a sphere and measured the size of the screen, distance of the eye to the screen, the angle of the eye center relative to the plane that the screen lay in, and the position of the eye within the screen. Using this information, we were able to map pixels to their corresponding visual degrees. We added an indicator that was synced to the presentation of each stimulus and detected via a photodiode in order to sync our stimulus to our electrophysiological recordings. For stimulus presentation, we used the PsychoPy package [43]. Stimuli were displayed using a Texas Instrument Lightcrafter PRO4500 in monochrome mode (green) running at 180Hz. The mean luminance of the projector was 1.39 *W/m*^2^, while the max luminance was 4.37 *W/m*^2^.

- White noise stimulus: (Figure 2, Figure 4) our white noise stimulus consisted of 5° horizontal bars flickering at 60 Hz with luminance values randomly drawn from a truncated Gaussian distribution. The stimulus was therefore changing across one spatial dimension and one time dimension, allowing for the extraction of two-dimensional spatiotemporal filters via white noise reverse correlation.
- Full field flashes: (Figure 3, Figure 4 Figure S5) OFF flashes of 20 ms, 40 ms, 80 ms and 160 ms with 10 second intervals were repeated for four sweeps per recording. High contrast OFF flashes consisted of light decrements from the mean luminance of the projector to its minimum output, corresponding to a Weber contrast of -1 (Figure 3A, Figure 4E), while low contrast OFF flashes consisted of light decrements from the mean luminance of the projector corresponding to a Weber contrast of -0.1.
- Ternary noise: (Figure S4) The ternary noise stimulus consisted of 5° horizontal bars flickering at 60Hz with luminance values randomly sampled from Weber contrast steps of -1, 0, or 1 (high contrast condition) from the mean luminance of the projector, or -0.1, 0, or 0.1 (low contrast condition) from the mean luminance of the projector.
- Drifting gratings: (Figure S8) Drifting grating stimulus consisted of 0.5 Hz, high contrast drifting square waves of spatial wavelengths ranging from 2.5°, 10°, 12.5°, 25°, 40°, 50°, 80°, 100°, 125°, and 200°.

### Reverse correlation for extraction of white noise filters

We extracted spatiotemporal white noise filters and static nonlinearities via the reverse correlation method as described in Behnia et al. [24] and elsewhere [23, 40, 4, 5]. All “white-noise filter” predictions in this study are linear-nonlinear (LN) predictions, as cell response predictions combine white noise (linear) filters with static nonlinearities.

To extract white noise filters for each cell, we selected continuous responses to white noise over a window of time ranging from 30-120 seconds depending on recording stability. Traces were downsampled to 100 Hz, and filters were extracted for a duration of 5 seconds. Spatiotemporal filter properties were not significantly affected by different downsampling factors, or by increasing or decreasing filter duration.

All spatiotemporal filters were space-time separable: thus, after a 2D spatiotemporal filter was extracted via reverse correlation, we extracted a characteristic 1D temporal filter by selecting the temporal trace at the spatial location with the highest amplitude. These 1D temporal filters were averaged across individual recordings to get a characteristic temporal filter for each cell type (Figure 2A, 4A). In order to characterize each temporal filter in frequency space, we convolved each 1D temporal filter with 1D sine waves of varying temporal frequencies from 0.1 to 10 Hz. The maximum steady-state amplitude of the convolved response at each frequency constituted a frequency tuning curve. These tuning curves were normalized and averaged across individual recordings to get a characteristic frequency tuning curve for each cell type (Figure 2B, 4B).

We extracted a characteristic 1D spatial receptive field by selecting the spatial profile at the time point with the highest amplitude. These 1D spatial receptive fields were averaged across individual recordings to get a characteristic spatial filter for each cell type (Figure 2C, 4C). As the white noise stimulus consisted of 5° horizontal bars, these spatial receptive fields have a resolution of 5°.

In order to obtain static nonlinearities, 2D white noise filters were convolved in time and summed in space to obtain (1D) linear predictions in time that could be compared with the (1D) recorded responses. The predicted and actual responses were binned by amplitude and averaged within each bin across recordings (Figure 2D, 4D). Bin size did not significantly affect static nonlinearity shape.

In order to compare flash responses to predictions based on extracted white noise filters, each spatiotemporal white noise filter was convolved in time with a 2D flash stimulus of the appropriate duration and summed across space. The resulting 1D linear prediction in time was then transformed via the static nonlinearity, resulting in a LN prediction. These LN predictions were then averaged (Figure 3A, S5A). The same approach was used to compare drifting grating data with white noise filter predictions (Figure S8).

## Parameter Fitting

We parameterized both extracted white noise filters and flash responses in order to compare Tm cell changes across conditions.

### Parameterization of White Noise Filters

Spatial receptive fields in all scenarios were fit to a Gaussian function *g*(*x*) = *e*^*−*(*x−µ*)^2^*/*2*σ*^2^^. The mean temporal filters for Tm1, Tm2, Tm4 and Tm9 were similarly fit with a biphasic function in time *t*:

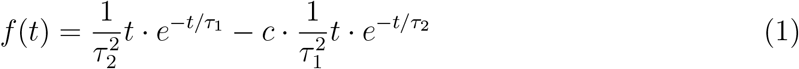

The two lobes of the biphasic function are determined by constants *τ*_1_ and *τ*_2_. For parameterizing temporal filters from our white noise analysis, we set *c* = 1. This constrained the convolution of the above function with a constant stimulus to integrate to zero, thus fitting the band-pass character of recorded cells (see Figure S2A,B). These parameterizations did not adversely affect the tuning properties of the filters for each cell type (Figure S6). For parameterized flash responses, *c* was unconstrained. All functions were parameterized using `scipy.optimize.curve fit.`

We derived frequency tuning curves for parameterized white noise filters by convolving them with 1D sine waves of varying temporal frequencies varying from 0.1 to 10 Hz. The tuning curve consisted of the maximum amplitude of the steady state response at each frequency (Figure 5C). These frequency tuning curves were identical to tuning curves derived analytically via transfer functions (not shown). The full width half max (FWHM) and peak frequency was calculated numerically (Figure 5D). To compare flashes with white noise filters in the same parameter regime, we generated white noise filter LN predictions of 20 ms flashes (Figure 5A,B) and plotted them alongside parameterized 20 ms flash responses.

## Linear Regression

In order to determine if our electrophysiological recordings of Tm1, Tm2, Tm4 and Tm9 could match electrophysiological recordings of T5, we applied linear regression of Tm1 and Tm9 flash responses to recorded T5 responses from Gruntman et al. [8]. The authors of this paper recorded individual T5 cell responses to static vertical bar flashes of width 2.25°, 4.5° and 9° at different spatial locations, and for a duration of 40 ms and 160 ms, for a total of six conditions. T5 Traces from Gruntman et al. [8] were accessed via https://figshare.com/collections/Simple_integration_paper_data_and_code/3955843.

We required coefficients to be strictly positive so as to maintain the sign of the input, and also did not fit an intercept under the assumption that all T5 recordings were preprocessed such that they had a baseline of zero. Regression was done using the `scikitlearn` LASSO module, which allows positive weight constraints, with α = 0.0001 (α = 0 equivalent to a simple linear regression). We first applied linear regression to the average T5 responses for each bar location and condition (Figure 6A-C, S7). We then applied linear regression to individual T5 traces for each T5 cell, for each bar location and condition (Figure 6D-F).

As input to the linear regression, we used: (1) Tm1 and Tm9 white noise LN predictions for 40 ms and 160 ms flashes, as well as (2) measured Tm1 and Tm9 response to 40 ms and 160 ms flashes, and (3) measured Tm1 and Tm9 response to 40 ms and 160 ms flashes in the presence of OA (Figure 6A,B). None of these inputs were parameterized.

Since our linear regression did not use an intercept term, we used the square of the sample Pearson correlation coefficient *r*^2^ as our measure of goodness of fit, instead of the coefficient of determination *R*^2^ [44]. *r*^2^ values were averaged across spatial locations for each condition and linear regression fit (Figure 6B).

Gruntman et al. [8] also recorded T5 responses to moving bars consisting of 20, 40, 80 and 160 ms consecutive flashes, across 2.25°, 4.5° and 9° widths. In order to predict the T5 response to moving bars, we summed the weighted Tm1 and Tm9 flash responses with appropriate time delays for the preferred direction and (opposite) null direction. The regression coefficients fit to the static T5 data were used for each matching condition (e.g. the coefficients from the 160 ms, 9° static condition were used to predict the response to the 160 ms, 9° moving bar condition, etc.). Both the PD and ND summed traces were then scaled by a single “gain factor” obtained by a separate linear regression on the combined PD and ND traces (Figure S7D).

## Model Construction

We built our framework for T5 based on established EM connectivity and an assumption of positivity for all Tm1, Tm2, Tm4 and Tm9 inputs onto T5. Specifically, Tm1, Tm2 and Tm4 were centered and Tm9 was offset by Δ*x* = 5° [45, 4]. The output of each of these cells was assigned a positive (cholinergic) connection weight proportional to EM synapse counts before being summed (Figure 7A, see below).

In order to construct a white noise model of T5 based on LN predictions for each cell type Tm1, Tm2, Tm4 and Tm9, 2D spatiotemporal receptive fields for each cell were constructed by taking the outer product of the parameterized gaussian spatial receptive field *g*(*x*) and the temporal filter *f* (*t*):

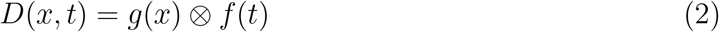

A given 2D stimulus in space-time *S*(*x, t*) is convolved with each spatiotemporal receptive field in time (but not in space), and then summed over space to give a 1D time course for each cell Tm1, Tm2, Tm4, Tm9. In discrete time this is:

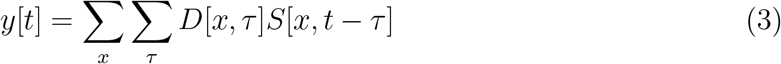

Finally, the mean of the static nonlinearities extracted via white noise analysis for each cell were parameterized by a softplus function:

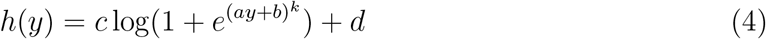

where *a* determines the sharpness of the “bend,” *b* translates the softplus curve along the x-axis, the multiplicative factor *c* controls the angle/slope, *d* determines offset along the y-axis, and the exponent *k* increases the curvature. The LN output of each cell was then normalized based on the numerical frequency tuning curve (so that the maximum possible gain across all frequencies was 1). Finally, Tm1, Tm2, Tm4 and Tm9 were scaled in a relative manner determined by the ratio of synapse counts from EM connectome data (see below) [18].

In order to construct a flash model of T5 based on the flash responses of Tm1, Tm2, Tm4 and Tm9, we parameterized responses to 20, 40, 80 and 160 ms flashes and constructed spatiotemporal receptive fields by taking the outer product with parameterized spatial receptive fields derived from white noise spatial filters with a spatial resolution of 2.25°. In order to simulate responses to moving bars, we summed temporal responses at each location with appropriate temporal delays for the PD and ND directions. We did not explicitly model bar width (as we had Tm responses to full field flashes but not to different bar widths), hence the predictions for each model in Figure 7E are the same across the x-axis. Like the white noise model, relative scaling between Tm1, Tm2, Tm4 and Tm9 was determined by the ratio of synapse counts from connectome data (see below) [18]. Spatial receptive fields were those extracted from white noise. We did not include static nonlinearities, as our recorded flash responses already represent the nonlinear processing properties of each cell.

## Direction Selectivity Index

In order to match measurements of direction selectivity between our model output and those used in the T5 datasets, we use two metrics that we call DSI_max_ and DSI_mag_.

Wienecke et al. 2018 [20], inspired by [46], use the “peak-to-trough” response to calculate DSI_mag_:

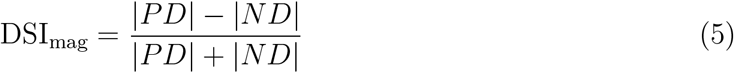

where |*PD*| represents the response magnitude to motion in the preferred direction, and response magnitude was calculated as 95th percentile minus 5th percentile. This works well to characterize steady-state responses to sine waves, and this metric is used in Figure 7C for *both* the Wienecke et al. [20] T5 sine wave data and the Gruntman et al. [8] T5 sine wave data. However, this measure is less amenable to transient flash responses. DSI_mag_ ASAP2f values (Figure 7C) were provided by Wienecke et al. [20]. DSI_mag_ values for T5 electrophysiology sine wave data from [8] were calculated using average peak and average trough values for both PD and ND traces.

Gruntman et al. [7, 8] use the following metric to describe their flash responses:

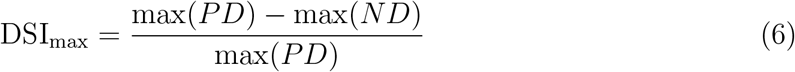

where each response max is defined as the 0.995 quantile within the stimulus presentation window. However, this does not take into account the ND amplitude in the denomenator, and is possibly susceptible to spuriously large DSI values due to noise [46]. We therefore use the following DSI_max_ for flash responses:

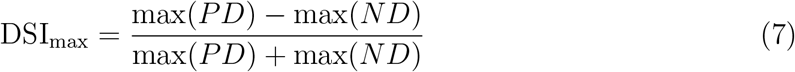

## Connectome Data

T5 synapse-level connectomic data was accessed from the comprehensive electron-microscopy (EM) reconstruction of inputs to T4 and T5 cells in the *Drosophila* optic lobe by Shinomiya et al. [18]. Detailed data from twenty reconstructed T5 cells is available, with synapse counts for each presynaptic cell Tm1, Tm2, Tm4, and Tm9 from various columns (https://flyem.dvid.io/fib19-grayscale). For a given T5 cell, we summed the synapse counts for each input (e.g. the synapse counts of Tm9 from column “K” and Tm9 column “C” were summed) and calculated the relative ratio of each of the four cell types. As reported in the study, Tm9 cells were consistently clustered on the leading edge of a given T5 cell, while Tm1, Tm2 and Tm4 cell synapses were clustered in the center of T5 dendrites. We therefore made the reasonable assumption that all synapse counts for each cell from various columns should be treated as a single offset (Tm9) or centered unit (Tm1,Tm2,Tm4). Twenty model instances were generated with these relative weight ratios, and the average PD tuning, ND tuning and DSI tuning were calculated (Figure 7B-C). The same approach was applied to flash models (Figure 7D-E). While a wide range of relative weight combinations confer direction selectivity on T5, we found that EM-based synaptic counts provide good fits across multiple models, suggesting that they are a reasonable estimation of synaptic weights in this system.

**Figure S1:**
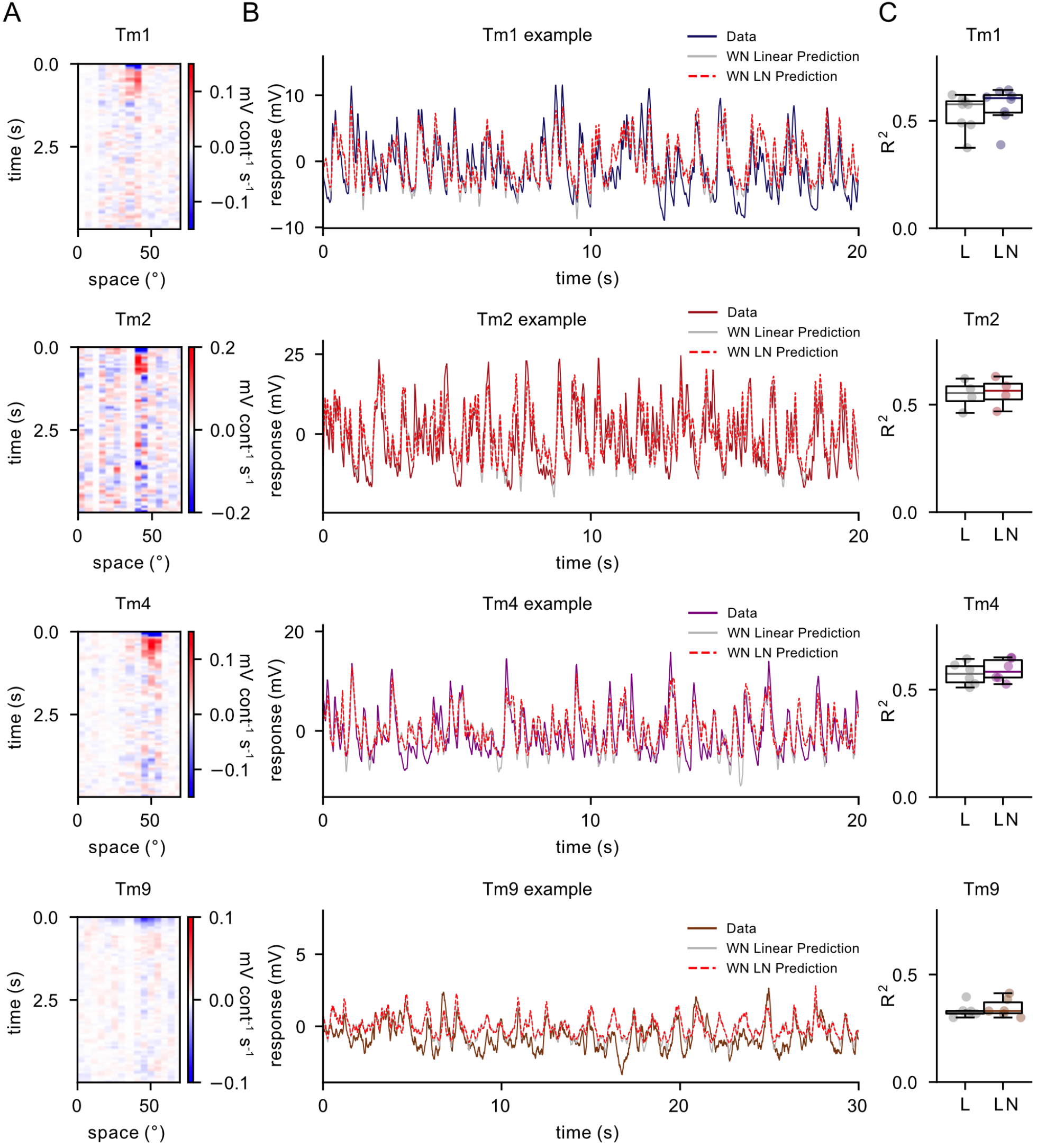
White Noise Analysis. **A**. Example white noise spatiotemporal linear filters extracted for single Tm1, Tm2, Tm4 and Tm9 neurons. **B**. Comparison of raw data (colored line), white noise filter linear prediction (grey line) and the white noise filter linear-nonlinear (LN) prediction (dashed red line) for the same neurons as in A. **C**. *R*^2^ values are comparable between linear and linear-nonlinear (LN) predictions for all cells.

**Figure S2:**
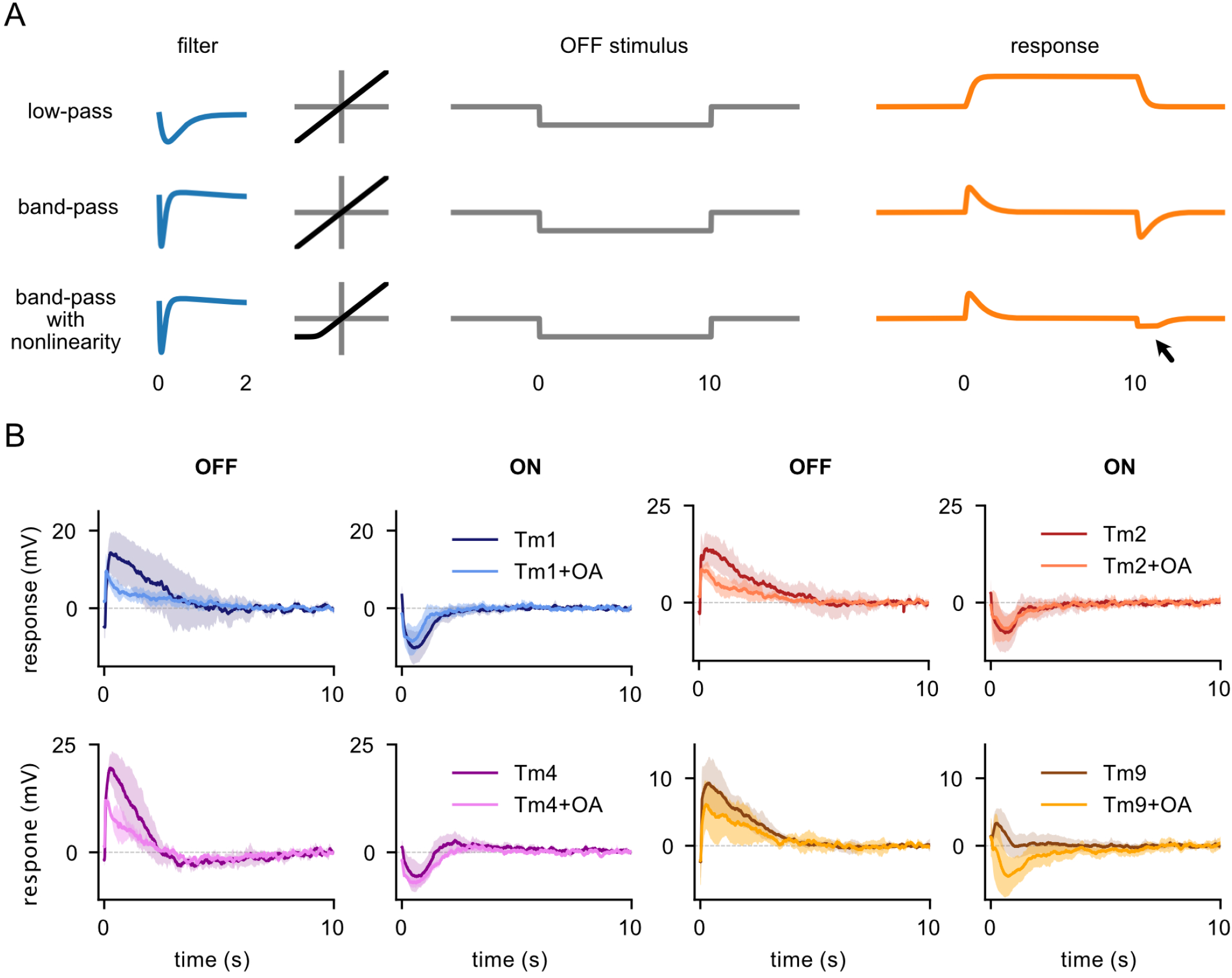
Long OFF and ON responses exhibit band-pass properties and are partially rectified. **A**. A low-pass filter (*top*) produces a response that fails to return to baseline until the stimulus ends, while a band-pass filter produces a response that returns to baseline during the course of a long stimulus. A linear band-pass filter produces symmetric responses to OFF and ON stimuli (*middle*), while a partially rectified band-pass filter produces asymmetric response to OFF and ON stimuli (*bottom*) **B**. Tm1 (n=4 saline, n=4 OA), Tm2 (n=6, n=4), Tm4 (n=4, n=2) and Tm9 (n=11, n=10) responses to 10 s OFF flashes and 10 s ON flashes in saline and in OA. All four neurons return to baseline during the flashes and therefore exhibit bandpass properties. They all show partial rectification in their ON responses. Tm9s presented more variability in their responses, with some cells showing depolarizing ON responses, resulting in depolarizing ON average.

**Figure S3:**
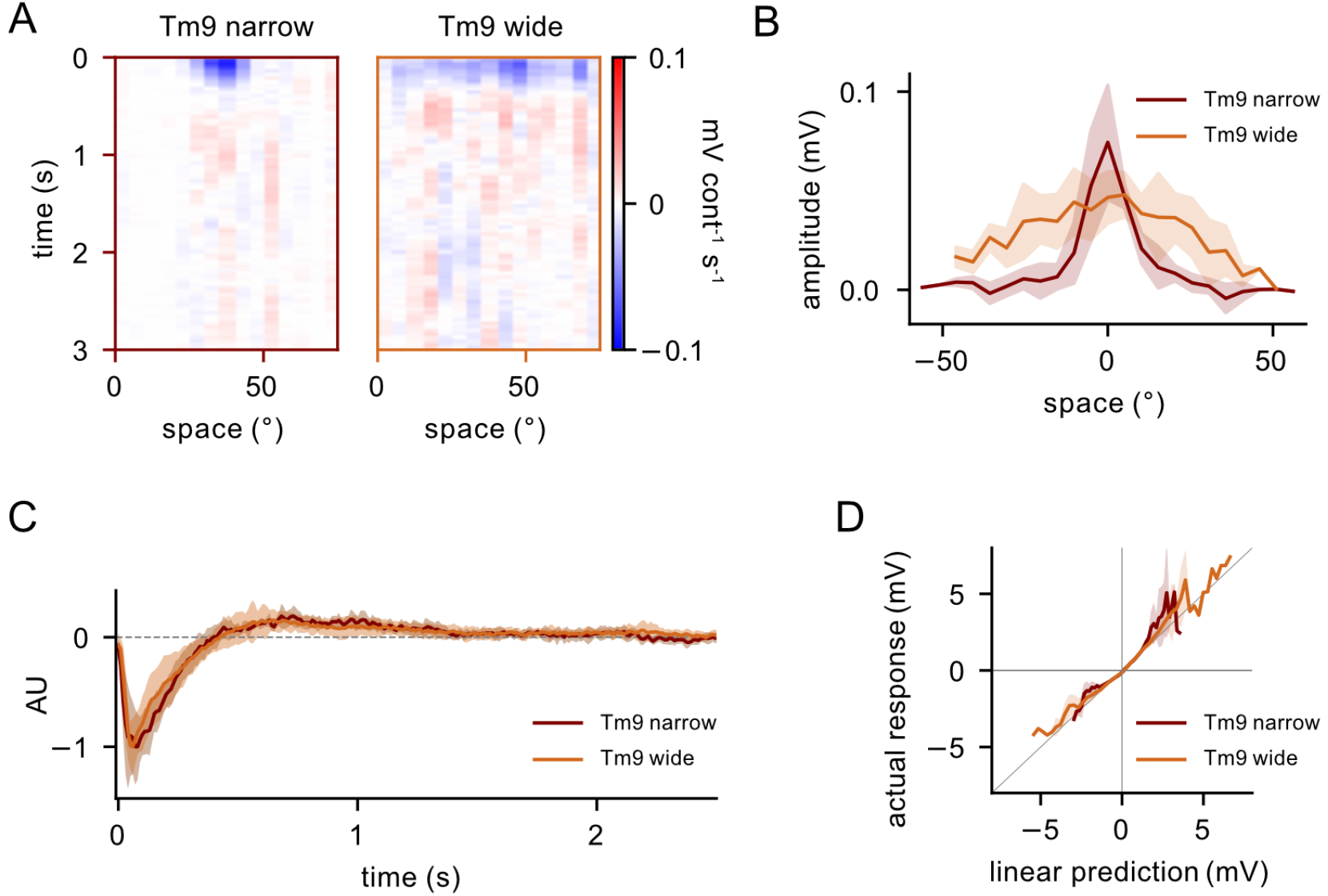
Tm9 cells fall into two clusters with narrow and wide spatial receptive fields. **A**. Example spatiotemporal receptive fields for narrow (n=6) and wide (n=8) Tm9 cells **B**. Narrow and wide spatial receptive fields (FWHM=15.4°, FWHM=60.3° when fit with a Gaussian) **C**. Temporal filters do not significantly differ **D**. Static nonlinearities do not significantly differ

**Figure S4:**
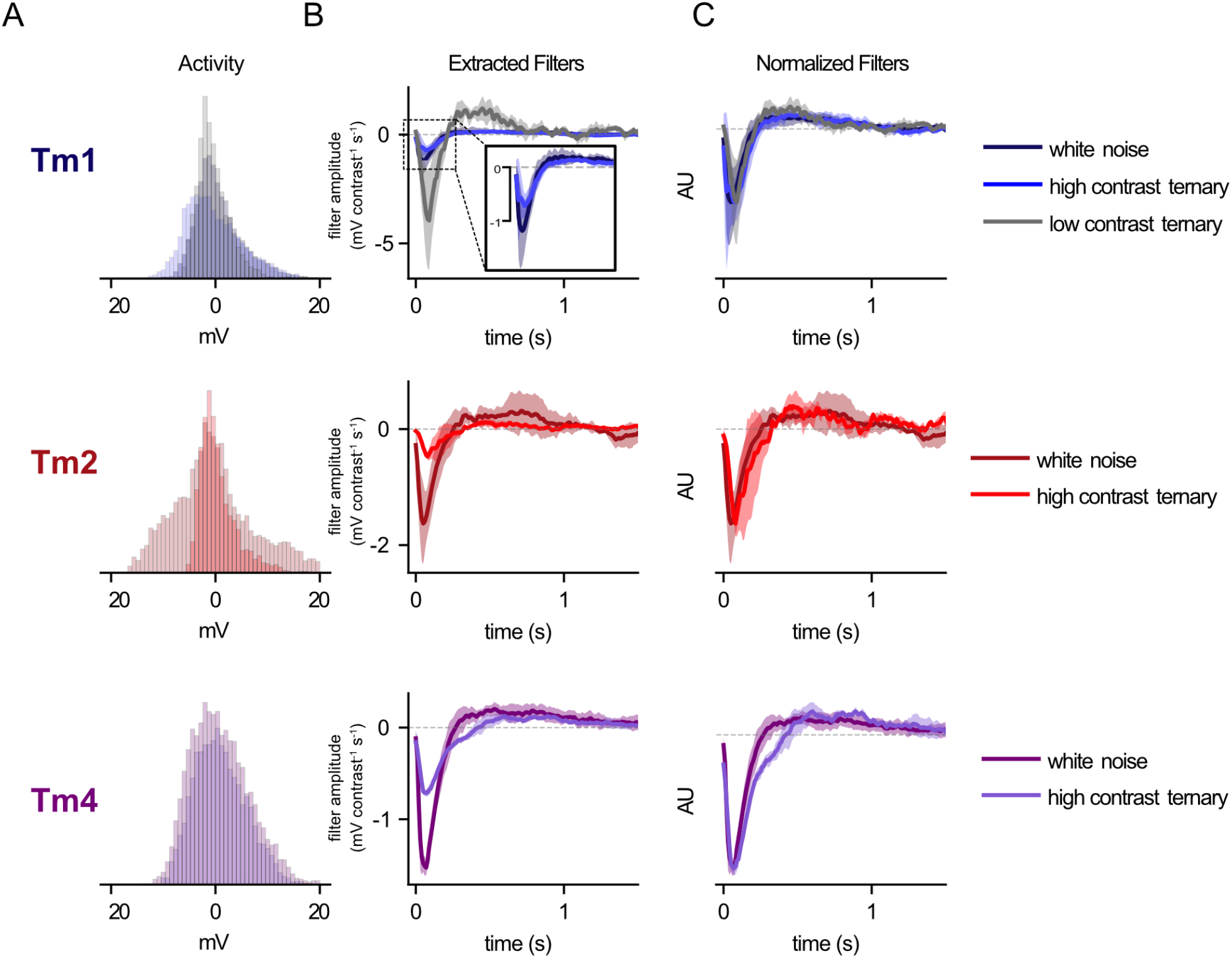
Tm1, Tm2 and Tm4 exhibit gain contrast adaptation. **A**. Histogram of response values across conditions. Different conditions elicit responses in the same general dynamic range, although low contrast responses overall have slightly lower amplitude **B**. Average temporal filters extracted from truncated white noise (*µ* = 0, *σ* = 1, truncated at *±*1), high contrast ternary noise (values randomly selected from +1, *−* 1 and 0) for Tm1 (n=4), Tm2 (n=2) and Tm4 (n=2), white noise (same data as in Fig. 2A), and low contrast ternary noise (values randomly selected from +0.1, 0.1 and 0) for Tm1 (n=4). Filter amplitude differences indicate gain adaptation so that response of cell is within similar dynamic range regardless of contrast. While this is especially evident in the case of the low contrast ternary noise-extracted Tm1 filter (*top*, grey trace), the same effect can be seen between high contrast ternary noise and the Gaussian white noise (lower contrast)-extracted filters for Tm1 (*inset*), Tm2, and Tm4. **C**. When scaled, filters do not show strong differences in shape

**Figure S5:**
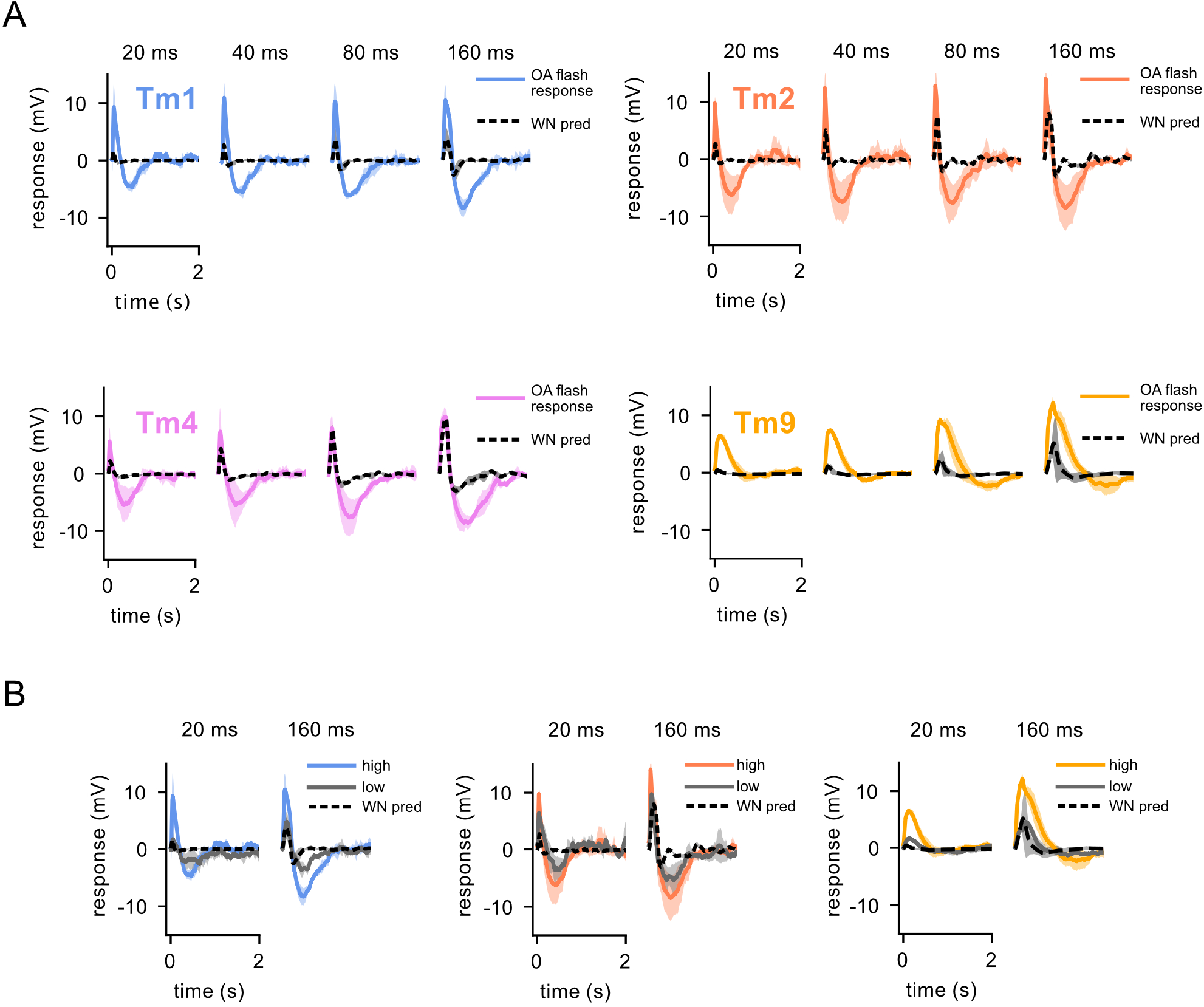
Tm1, Tm2, Tm4 and Tm9 flash responses in OA. **A**. Flash responses of T5 columnar inputs to 20, 40, 80, and 160 ms flashes in the presence of OA, with OA white noise filter predictions superimposed (black dashed line). Same data as in Figure 4E **B**. Tm1, Tm2, and Tm9 responses to flashes of high vs. low contrast (n=4, n=3, n=2, respectively) in the presence of OA. OA white noise filter predictions superimposed (black dashed line).

**Figure S6:**
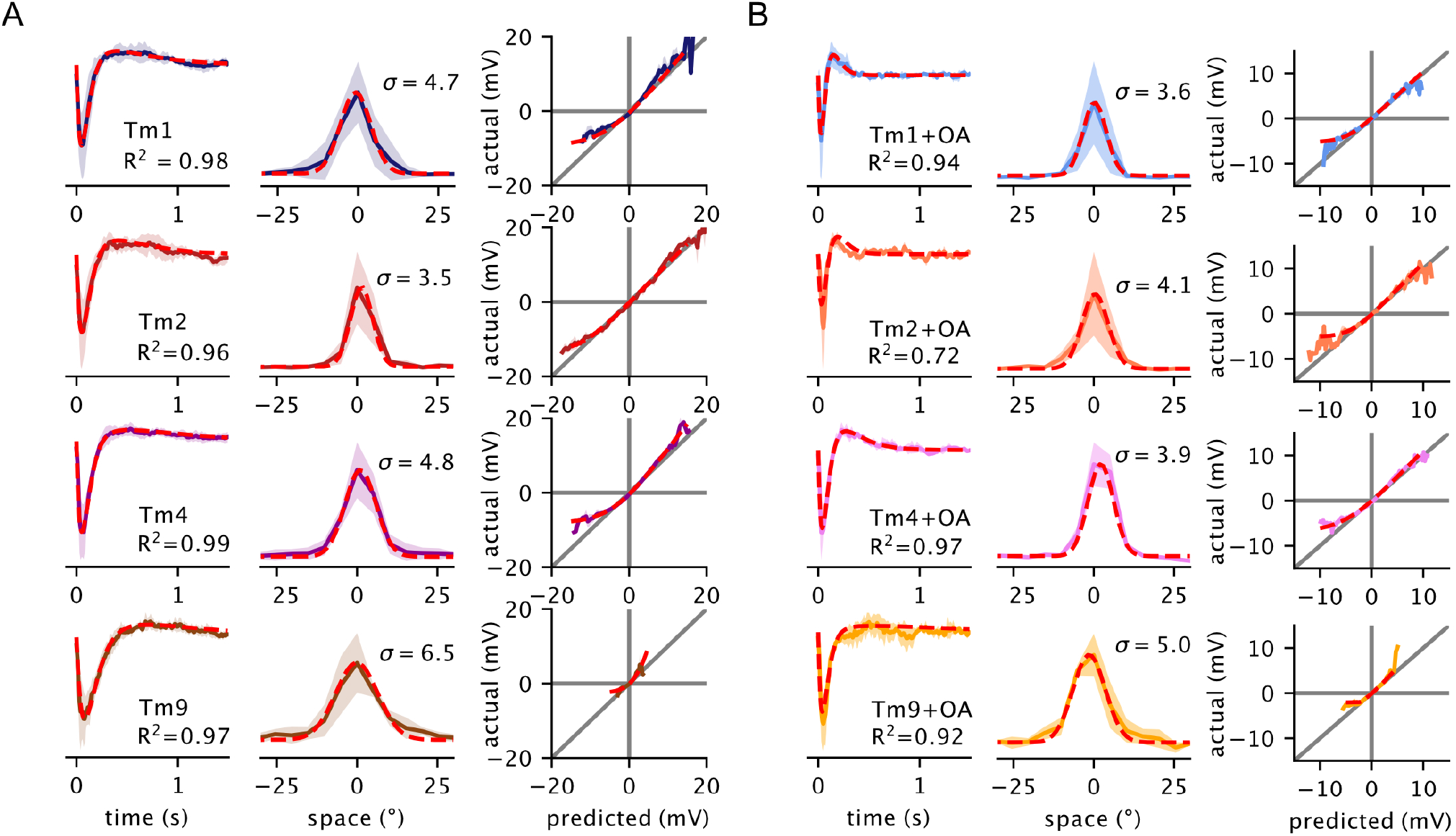
Paramaterization of white noise filters. **A**. Parameterization of white noise temporal (*left*), spatial filters (*middle*), and static nonlinearities (*right*). Same traces as in Figure 2, with parameterization superimposed (red dashed line) **B**. Same for OA, with traces from Figure 4.

**Figure S7:**
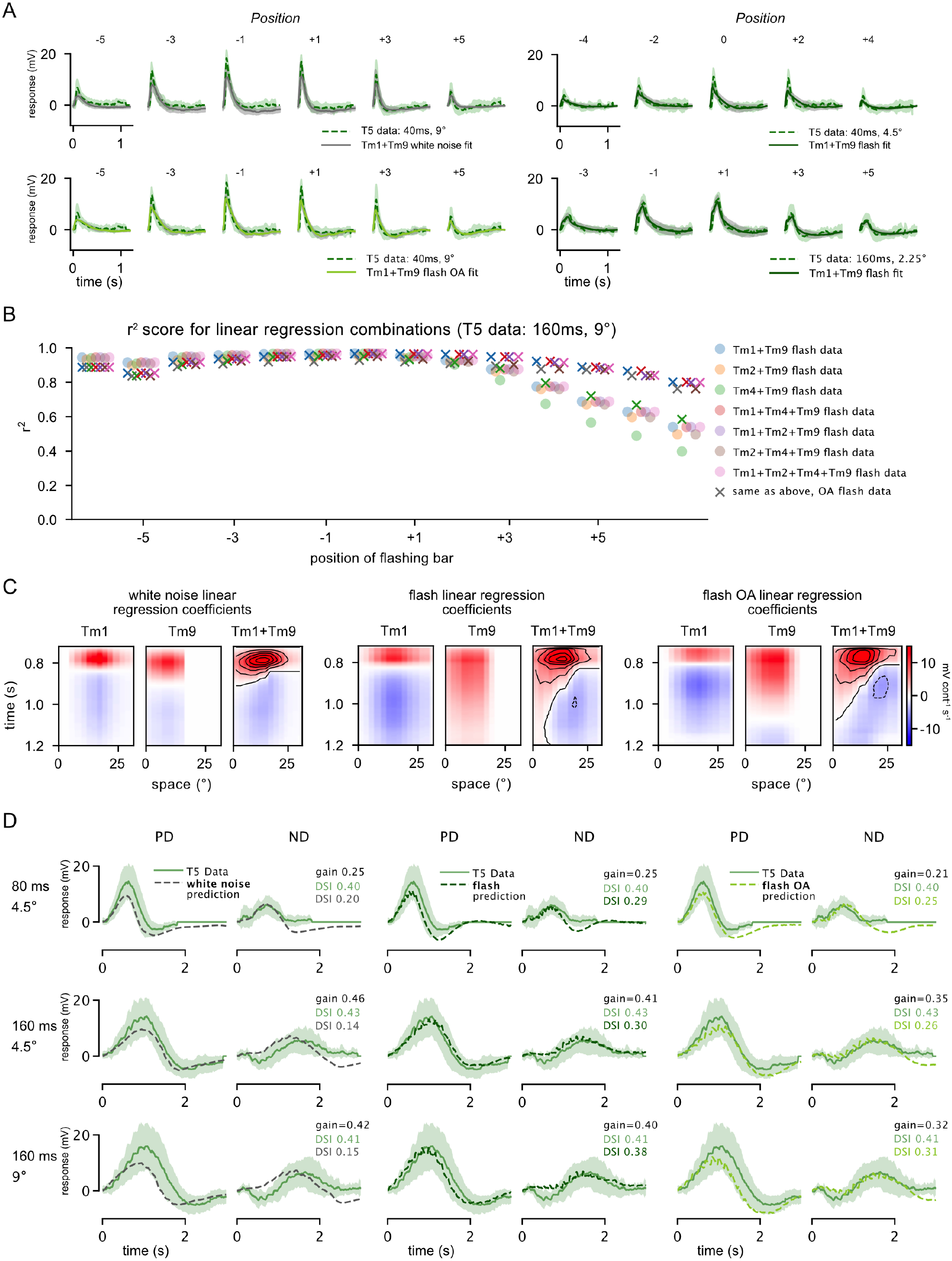
Extended linear regression analysis. **A**. Four examples of linear regressions from individual stimulus conditions varying in the length of stimulation (40 ms and 160 ms), as well as stimulus size (9°, 4.5*°*, and 2.25°). T5 data from Gruntman et al. [8]. **B**. In Figure 6, we chose to apply linear regression with Tm1 and Tm9. Combinations of Tm1, Tm2, and Tm4 with Tm9 perform approximately equally well (saline fits shown in circles, OA fits shown with crosses). **C**. Tm1 and Tm9 weighted by linear regression coefficients at each spatial location in the 160 ms, 9° condition, for the three fits enumerated in Figure 6 (using white noise filter predictions, flash responses, and flash OA responses, see Methods). The weighted Tm1 and Tm9 components are summed to generate a representative spatiotemporal receptive field (*right of each panel*). **D**. Gruntman et al. [8] recorded T5 responses to moving bars across multiple stimulus conditions (20, 40, 80, and 160 ms duration and 2.25°, 4.5*°*, and 9° bar width). Linear regression coefficients fit to static flashes across conditions (160 ms, 4.5° and 9°) predict T5 moving bar temporal responses (see Methods). In particular, the Tm1 and Tm9 flash data in the baseline saline condition and OA condition match the T5 electrophysiology traces, as well as DSI (*center column, right column*). Note that both PD and ND traces are scaled by a single “gain” factor (see Methods).

**Figure S8:**
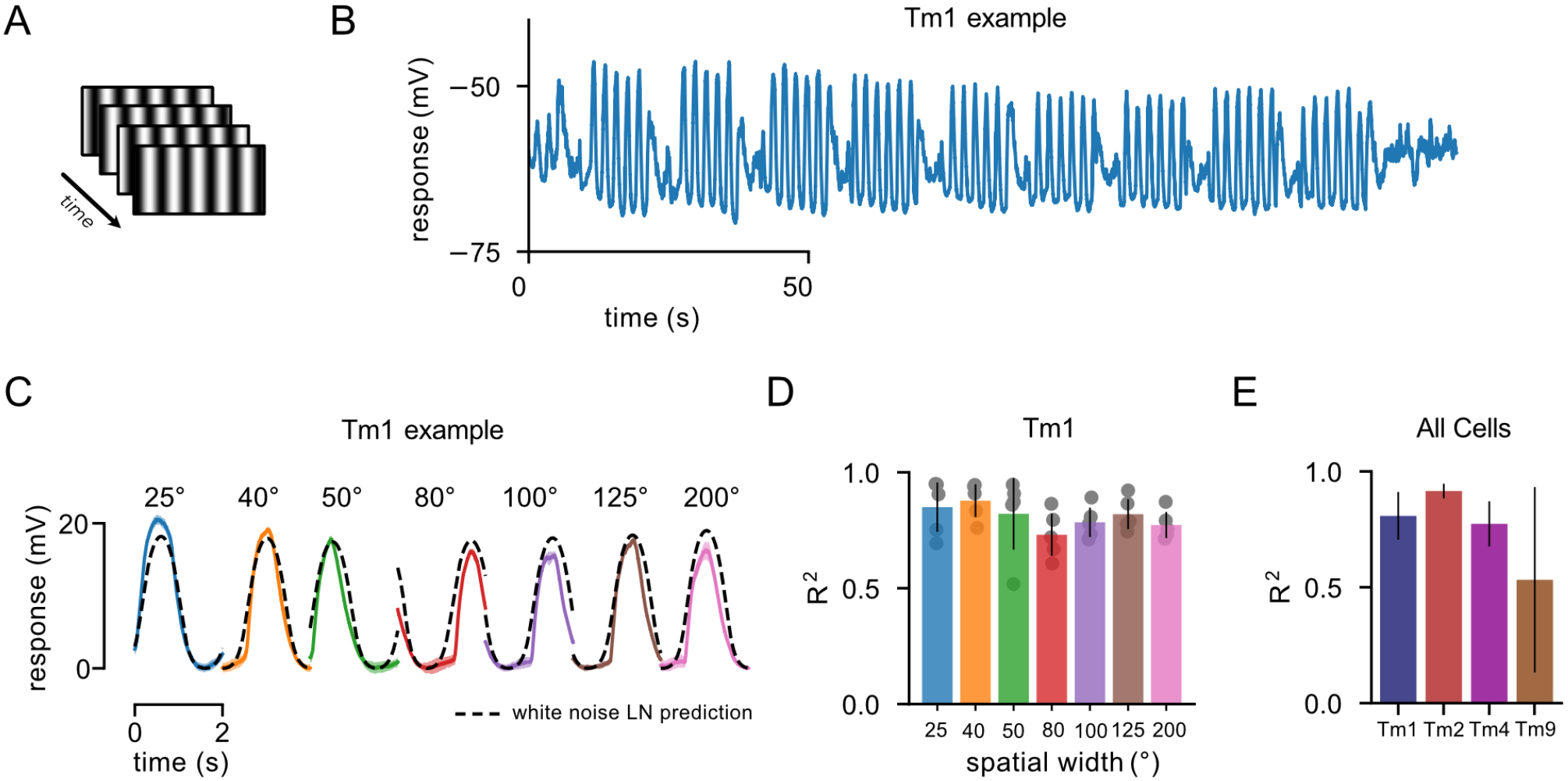
Drifting grating responses are well predicted by white noise filters. **A**. Drifting gratings with 0.5 Hz temporal frequency and with varying spatial frequencies were shown **B**. Raw drifting grating response for example Tm1 cell **C**. Averaged periodic responses for each spatial frequency (colored traces for a single example Tm1 cell). A linear-nonlinear prediction based on the corresponding spatiotemporal white noise filter captures the temporal aspects of the response (dashed black line). **D**. *R*^2^ responses for each stimulus condition across Tm1 cells (n=5). The match between predicted and actual responses indicates that using a white noise filter linear-nonlinear framework to model T5 responses to drifting gratings is reasonable. **E**. *R*^2^ responses for Tm1 (n=5), Tm2 (n=1), Tm4 (n=2) and Tm9 (n=2) averaged across stimulus conditions.

